# Heterogeneity in cilia patterning enables multiple flow functions in Paramecium

**DOI:** 10.64898/2026.02.19.706812

**Authors:** Daphne M. Laan, Aikaterini M. Kourkoulou, Guillermina R. Ramirez-San Juan

## Abstract

Free-living unicellular organisms known as ciliates rely on fluid flows to perform essential functions. These flows emerge from the coordinated activity of thousands of cilia organized into arrays with highly diverse architectures. Within a species, the structure of the ciliary array is highly stereotyped and persists in time, raising the question of how a single cell achieves multiple distinct flow functions from one fixed array architecture. Here, we investigate how the ciliary array in the ciliate *Paramecium tetraurelia* enables this organism to feed and swim simultaneously. Using expansion microscopy and high-speed imaging, we measure ciliary organization and kinematics, from individual beat dynamics to collective metachronal wave patterns, together with the flows these cilia generate. We find that the cell’s surface is partitioned into domains with distinct spatiotemporal cilia organization. In addition to the oral apparatus, there are two structurally different domains. One is a densely ciliated, high-frequency beating domain located anterior to the oral apparatus, and the other covers the remainder of the cell’s surface, where cilia are more sparsely distributed and slower-beating. Selective removal of each results in impaired feeding or swimming, respectively, demonstrating that each domain is functionally specialized. Our work establishes heterogeneity in spatiotemporal ciliary patterning as a mechanism that allows a single cell to generate multiple flow functions. By measuring cilia organization and the flows they generate, we uncover general physical principles by which cilia arrays produce flow patterns that underlie diverse biological functions.

---

Ciliates are unicellular eukaryotes, found in virtually all freshwater and marine environments, that exhibit remarkable diversity in morphology and lifestyle [1, 2]. A defining feature of these organisms is their highly patterned cortex, which is covered by dense arrays of motile cilia. The coordinated beating of these cilia generates macroscopic fluid flows underlying essential functions such as swimming, navigation, and feeding [3–7].

In multicellular ciliated organisms, such as marine larvae, the architecture of flows generated by their cilia can be modulated through changes in ciliary kinematics under neuronal control, allowing the same cilia array to perform different functions [8, 9]. However, in free-living unicellular organisms such as ciliates, it remains unclear how different flow functions are achieved within a single cell. Here we address this question in *Paramecium tetraurelia*, a ciliate that uptakes food particles while swimming [10–12]. In the absence of a neural system, any distinct flow functions this cell generates must instead be encoded in the structure and kinematics of its ciliary array.

*Paramecium tetraurelia* has long served as a model system for studying cilia, particularly in the context of basal body biogenesis, structure and duplication [13–15]. Previous studies have noted that the cell has cilia populations with different basal body organization, biochemical modifications, and beating patterns [16–18]. However, quantitative measurements of cilia and flow organization across the entire organism have been out of reach, because they require resolving individual basal bodies across the whole cell surface and capturing cilia fast dynamics. As a result, we lack an understanding of the relationship between the spatiotemporal organization of cilia and the resulting flow structure, which is essential for understanding how a single cell can generate multiple flow functions.

Modern experimental techniques such as expansion microscopy, high-speed imaging, and quantitative image analysis now make these measurements possible. Using these approaches, here we map cilia architecture and kinematics across scales over the entire surface of *Paramecium tetraurelia*. We quantify cilia organization across the cell and find that it is spatially partitioned into well-defined regions with distinct cilia array patterns and kinematics. Located anterior and to the right of the oral apparatus is a region where cilia are more densely packed and beat at a higher frequency than those in the remainder of the cell. Selective removal of these cilia results in loss of the flows required for feeding and an inability to ingest particles; remarkably, these cells do not display any defects in swimming. Conversely, cells retaining only the cilia anterior to the right of the oral apparatus, generate feeding flows and thus are able to ingest particles normally but are unable to swim.

Together, these results reveal that partitioning the cell’s surface into domains with distinct cilia spatiotemporal organization is the mechanism that allows unicellular organisms to perform diverse flow functions. More broadly, our work identifies and measures the key physical properties of a ciliary array that determine the flow patterns it generates and, consequently, its function. This provides an experimental structure-to-function map connecting cilium-level patterning (micron scale) to macroscopic flow function (swimming and feeding, millimeter scale). These results have implications for understanding the behavior and function of multiciliated cells across eukaryotes, and they uncover general design principles for engineering flows with specific functions at low Reynolds number.

## CILIA ARRAYS WITH DISTINCT DENSITY AND ORIENTATION ARE PATTERNED WITHIN A SINGLE CELL

*Paramecium* is an oblong ciliate with a length of 114.3±9.6 µm and a width of 40.3±3.5 µm (Fig. S1A). Its surface is completely covered by cilia that are arranged in highly stereotyped rows along the anterior–posterior (A–P) axis of the cell [19]. The *Paramecium* ciliary array is composed of the canonical structural elements found in ciliated eukaryotes. Each cilium is anchored to the cell membrane at its base by a basal body, linked to its neighbors by striated fibers, which are cytoskeletal structures that contribute to the anchoring and orientation of cilia [14, 20–24]. Differences in basal body number and ciliation across cortical regions of the cell have previously been noted [14]. However, we lack measurements of the physical parameters of a ciliary array that determine the flow structure it generates, such as cilia density, orientation, and collective beating patterns. Furthermore, how these parameters vary across the cell is unknown.

First, to measure the cellular-scale spatial organization of the *Paramecium* ciliary array, we used expansion microscopy (ExM), staining basal bodies and striated fibers (see Methods § 1). This technique allows us to resolve individual basal bodies and the orientation of single striated fibers across the entire cell. From basal body images, we observe on the ventral side of the cell the oral apparatus, which has at its opening a single densely packed row of cilia and occupies 3.5 ± 1.5% of the cell’s surface (Fig. S1C,E). Consistent with previous findings, basal bodies anterior to the oral apparatus occur in adjacent pairs, both ciliated, whereas in the rest of the ventral side, as well as across the dorsal side, basal bodies are found individually or, if adjacent, only the posterior basal body of each pair bears a cilium (Fig. 1A–B, S1B) [14, 19]. This difference in basal body pattern suggests a corresponding difference in cilia density across the array.

**FIG. 1.**
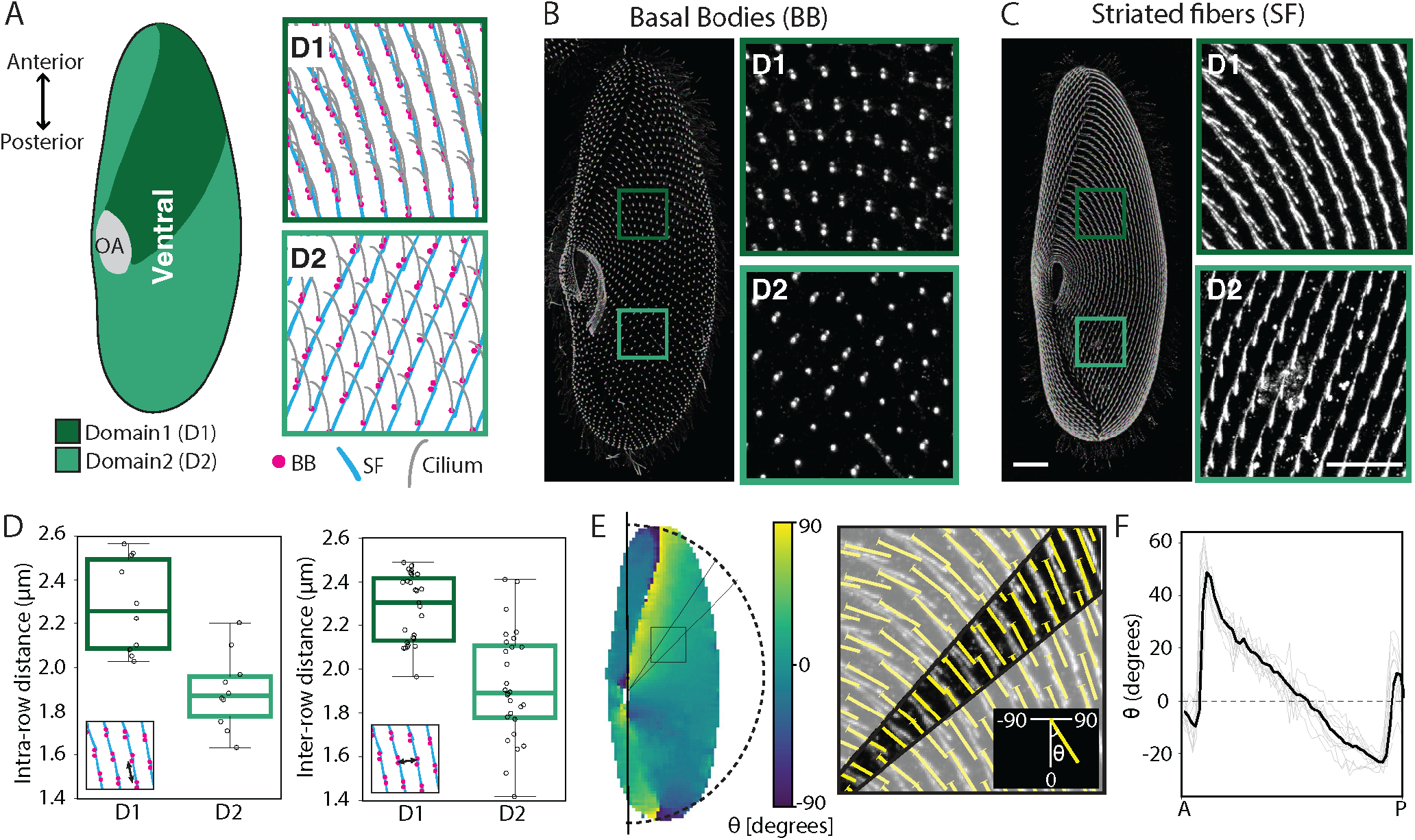
Highly organized cilia arrays with distinct architectures are patterned in the same cell. (A) Schematic of a *Paramecium tetraurelia* cell showing the anterior-posterior (A-P) axes and ventral cilia and basal body organization of the cell. Colors show the two domains with different cilia patterning, domain 1 (D1, dark green), domain 2 (D2, light green) and the oral apparatus (gray). Insets show schematics of the cilia, basal body (BB) and striated fiber (SF) organization in D1 and D2. (B-C) Maximum intensity projections of BBs and SF signal visualized by Expansion Microscopy (ExM). Ventral view of the cell is shown. Insets highlight the distinct organization in D1 and D2. The images in B and C are shown in the same scale. Scale bars: 10 µm and 5 µm in insets. (D) Basal body spacing in D1 and D2. Left: Intra-row distances between neighboring basal bodies or basal body pairs. Right: Inter-row distances between adjacent basal body rows. Mean intra-row spacing: D1 2.28 ± 0.22 µm, D2 1.89 ± 0.17 µm (n=10 cells). Mean inter-row spacing: D1 2.28 ± 0.16 µm (n=27 cells), D2 1.93 ± 0.25 µm (n=28 cells). Each point represents the average of one cell. The same cells were used for the measurements of the different regions. (E) Method for determining SF orientation. Left: Colormap of SF orientation angle *θ*, overlaid on the ventral cell surface. The cell was divided into 60 angular bands of 3° each, radiating from the oral apparatus and spanning the region between the anterior and posterior ends of the cell. SF orientation vectors were averaged within each sector. Inset: SF fibers in D1, overlaid with computed orientation vectors (yellow). Below: sign convention for *θ*, defined relative to the cell’s A–P axis (*θ* = 0°). (F) Average SF orientation profile on the ventral surface, computed across multiple cells as a function of position along the A–P axis. At each position, SF orientations were averaged within the corresponding angular sector (see E). Gray curves represent individual cells, and the black curve indicates the population mean (n=10 cells). Dashed line marks *θ* = 0° (fibers parallel to the A–P axis).

To test this, we compared the region anterior and to the right of the oral apparatus, where basal bodies occur in ciliated pairs, with the rest of the cell surface. We analyzed image tiles from two-dimensional projections of each region. To minimize projection artifacts, we selected tiles from the central part of each region, identified the center of each basal body, and measured the distance between them (see Methods § 2 a). In the anterior region, basal bodies within a pair are immediately adjacent, while successive pairs are separated by 2.28 ± 0.22 µm (Fig. 1D). In the rest of the cell, basal bodies are spaced more closely, with an average separation of 1.89 ± 0.17 µm (Fig. 1D). Despite this closer spacing, the lower fraction of ciliated basal bodies in the rest of the cell means that these differences in packing collectively result in an overall 1.4-fold increase in cilia density in the anterior region. We therefore term the denser region domain 1 (D1) and the less dense region domain 2 (D2).

To measure the surface area occupied by each domain, we took advantage of the fact that differences in cilia density are evident in our ExM images as differences in the intensity of the polyE channel. Thresholding this signal allowed us to calculate the area occupied by each of three distinct regions in the array (Fig. S1D Cilia; see Methods § 2 a). D1 lies anterior and to the right of the oral apparatus and occupies 16.1 ± 2.8% of the cell’s surface. D2 covers the rest of the ventral side together with the dorsal side and occupies 80.4 ± 3.0% of the cell’s surface. The oral apparatus (OA), located on the ventral surface, occupies the remainder of the cell’s surface (Fig. 1A, S1D,E).

Because the orientation of striated fibers (SFs) is expected to mark the local direction of ciliary beating [25– 27], we next quantified the organization of the SFs across the cell. On the ventral side, SFs form continuous rows that curve around the oral apparatus (Fig. 1C), whereas on the dorsal side they form rows parallel to the anterior– posterior (A–P) axis (Fig. S1B). To quantify their orientation, we used the alignment by Fourier transform workflow, which exploits 2D fast Fourier transforms to obtain the orientation of fibrillar features in microscopy images (Methods § 2, striated fiber orientation) [28]. This analysis yields a director field that describes the local alignment of the SFs (yellow lines, Fig. 1E). Next we measured the angle of each director relative to the A–P axis of the cell. Given the cell’s symmetry, we aligned this axis with the oral apparatus (Fig. 1E). To measure the variations of SF angle along the A-P axis of the cell we averaged the local orientation over radial bins (Fig. 1E, Methods § 2 a). We found that on the ventral side, SFs are oriented at an angle of ∼ 50° with respect to the A-P axis at the anterior end of the cell where D1 starts. Moving posteriorly, this angle decreases gradually until SFs become parallel to the axis (0°) approximately halfway along the cell, then continues to change smoothly, becoming negative as the SFs curve toward the opposite side, reaching a maximum deviation of ∼ 25° (Fig. 1F). In contrast, dorsal SFs remain largely parallel to the A–P axis, only deviating by less than 20° from it at the poles of the cell (Fig. S1F).

Together, these measurements show that the surface of *Paramecium* is partitioned into domains with distinct cilia density and SF orientation. Because SF orientation marks the direction of ciliary beating in multiciliated cells [26, 27], and collective beating patterns such as metachronal waves are sensitive to both ciliary spacing and orientation [29–31], these architectural differences between D1 and D2 suggest corresponding differences in ciliary kinematics. Therefore, we next measured individual and collective ciliary beating patterns across the cell.

## INDIVIDUAL CILIA DYNAMICS AND COLLECTIVE METACHRONAL WAVE PATTERNS VARY ACROSS DOMAINS

To generate fluid flows that support diverse biological functions, cilia arrays must coordinate the activity of individual cilia across entire surfaces. Individual cilia generate flow by beating with an asymmetric periodic stroke that can be decomposed into an effective stroke, which propels fluid, and a recovery stroke, during which the cilium bends closer to the cell surface and returns to its starting position. Neighboring cilia beat with a constant phase shift relative to one another, giving rise to collective beating patterns known as metachronal waves (Fig. 2A) [32]. Theoretical work has shown that, in addition to the spatial organization of cilia, the emergence of metachronal waves and their properties is determined by individual cilium properties such as beat frequency and the direction of the effective stroke [33, 34]. However, we lack experimental measurements that simultaneously measure individual cilia beat kinematics and metachronal wave properties of an entire organism. In *Paramecium*, previous work has measured beat frequency [17, 35], without measuring beat orientation and its spatial variation across the cell. Classical work has been limited to schematic depictions of metachronal waves derived from qualitative visual inspection, with wavelength inferred from fixed images of the cell boundary [36, 37]. Modern imaging techniques now enable live imaging of cilia arrays, and quantitative image analysis frameworks allow unbiased, direct measurement of individual and collective cilia kinematics from high-speed microscopy recordings [3, 38, 39].

**FIG. 2.**
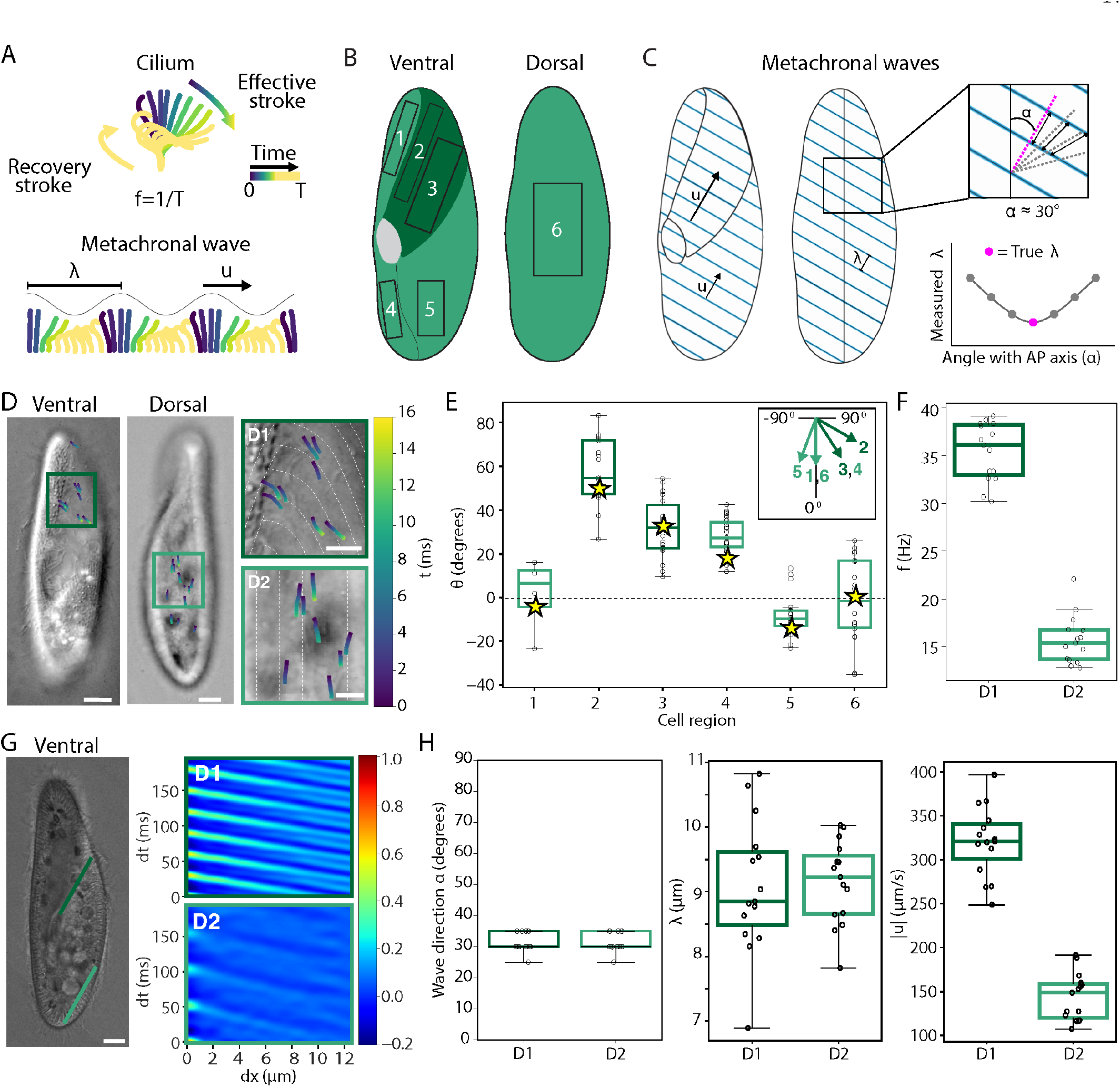
Cilia beat orientation and frequency varies across cell surface regions. (A) Top: Schematic of the ciliary beat cycle, consisting of an effective stroke (blue to green) that drives fluid flow and a recovery stroke (yellow). Bottom: Schematic illustrating coordination of neighboring cilia into a metachronal wave pattern, characterized by its wavelength (*λ*) and wave velocity (u). (B) Schematic of ventral and dorsal views of the cell. For the cilia beat direction measurements we divide the cell into 6 different regions. Regions 2 and 3 fall within D1, while regions 1, 4, 5, and 6 fall within D2. (C) Left: Schematic of ventral and dorsal views of the cell showing metachronal wave crests (blue) and propagation direction (arrows, length scales with wave velocity) in both regions. Both D1 and D2 have a similar wavelength (*λ*) and propagation direction (arrow), but different wave velocity (u). Right: Determination of the metachronal wave propagation angle (*α*) and wavelength (*λ*). Top: Inset shows the A-P axis and wave direction *α*. Bottom: Apparent *λ*, defined as the perpendicular distance between crests, measured from kymographs at increasing scan angles relative to the A-P axis. The true *λ* (magenta) corresponds to the minimum of this curve. (D) Representative frames from high-speed live DIC imaging of ventral and dorsal cell surfaces, overlaid with traces of ciliary tip trajectories during the effective stroke. Imaging was focused ∼ 10*µm* above the cell surface to enable tracking of ciliary tips. Insets include the SF direction, visible in white dashed lines. Scale bars: 10 µm and 5 µm in insets. (E) Orientation of the ciliary effective stroke relative to the A-P axis in different cell regions shown in B. Each point represents one cilium (n = 4, 17, 22, 23, 21, 18 cilia from 1, 4, 5, 5, 4, 4 cells for regions 1-6, respectively). The yellow star marks the average SF direction in the corresponding region (Fig. 1E-F, Fig. S1F). Top right: Reference frame used for all reported angles, with arrows indicating the average cilia beat direction *θ* per region. (F) Comparison of cilia beat frequency between D1 and D2 (n=15 cells). (G) Left: High-speed live DIC image frame of the ventral surface, with lines indicating the direction of wave propagation for both regions used to extract intensity profiles over time. Right: space-time autocorrelation of the intensity profile extracted on the line, showing global cilia coordination in both regions. Scale bar: 10 µm. (H) Comparison of metachronal wave parameters, wave direction, wavelength, and wave velocity, between D1 and D2 (n=15 cells).

To measure cilia kinematics with high spatiotemporal resolution, we immobilized cells by adhesion to a coverslip, and imaged them using ultra-fast DIC microscopy. First we characterized the individual beat direction of cilia (see Methods § 1 e and § 2 b). To determine the orientation and direction of the ciliary effective stroke, we tracked the motion of ciliary tips by imaging at a focal plane approximately 10 µm above the cell surface (Fig. 2D, Movie 1). At this height, only the part of the ciliary beat cycle where the cilium is most extended is captured, allowing selective visualization of the effective stroke. We found that in both D1 and D2 the effective stroke is directed towards the posterior side of the cell, while its trajectory follows the orientation of the underlying striated fibers (Fig. 2B,D-E). The beat direction is defined as the signed angle *θ* between the stroke trajectory and the A–P axis, where *θ* = 0° corresponds to a stroke directed towards the posterior pole (Fig. 2E, inset, which also shows the average beat direction in each cell region). On the dorsal side, the cilia effective stroke is parallel to the A-P axis, matching the SF pattern (Fig. S1F). On the ventral surface, *θ* varies with the average local orientation of the SFs, being greatest in D1 and smallest in D2 (Fig. 1E-F, Fig. 2E). Measurements of the direction of ciliary beat in the paroral membrane, which surrounds the opening of the oral apparatus, show that oral cilia beat toward the opening of the oral apparatus with an effective stroke oriented at 80.6 ± 7.7°, nearly perpendicular to the A-P axis (Fig. S2A, Movie 1).

In both D1 and D2, we observe MWs that propagate robustly towards the anterior pole of the cell, opposite to the direction of the ciliary effective stroke (Fig. 2C, Movie 2). To determine the direction and properties of the MWs, we analyzed spatio-temporal patterns of ciliary motion within defined regions of the cell surface. In a metachronal wave, neighboring cilia beat with a constant phase difference, which gives rise to wave fronts, lines along which cilia beat in phase. The wave fronts are perpendicular to the direction of propagation, and the wavelength is the distance between successive fronts. The wavelength measured along a line is therefore smallest when the line is perpendicular to the wave fronts, that is, parallel to the direction of propagation. Along any other orientation, the line crosses the fronts obliquely and the apparent wavelength is larger (Fig. 2C). To find the direction of propagation in each region, we placed a line of interest along the direction of propagation observed in the recording and rotated it in both directions. From each line, we computed the two-dimensional autocorrelation of its kymograph and obtained the spatial wavelength from the dominant spatial peak (Fig. 2G). The orientation that minimized the wavelength was taken as the direction of propagation. We find that waves propagate at an angle of ∼ 30° relative to the A–P axis in both regions (Fig. 2H, Movie 2). The same autocorrelation analysis yields the parameters that describe ciliary kinematics, including beat frequency, metachronal wavelength, and wave speed (Fig. 2F-H; see Methods § 2 c).

We find that cilia beat frequency differs markedly between regions. Cilia in D1 beat at an average frequency of 35.4 ± 3.0 Hz, approximately twice that of D2 (15.7 ± 2.5 Hz) (Fig. 2F), consistent with the range reported in previous studies [17, 35]. Despite this frequency difference, the metachronal wavelength is nearly identical in both regions, measuring 9.1 ± 1.0 µm in D1 and 9.1 ± 0.6 µm in D2. Consequently, MWs travel approximately twice as fast in D1, with a velocity of 320.6 ± 39.9 µm/s compared with 143.9 ± 26.7 µm/s in D2 (Fig. 2H). Cilia forming the paroral membrane at the opening of the oral apparatus beat at 30.2 ± 2.1 Hz, close to the D1 frequency (Fig. S2B, Movie 2).

The relative orientation between the direction of MW propagation and the direction of the effective ciliary stroke is described as dexioplectic when the wave propagates perpendicular to the stroke, and antiplectic when it propagates opposite to it [40]. Because the wave propagates at a constant angle of ∼ 30° from the A-P axis in both domains (Fig. 2H), while the effective stroke direction varies across regions (Fig. 2E), the relationship between wave and stroke direction differs between domains. MWs are dexioplectic in D1, and both dexioplectic and antiplectic in D2.

Together, these measurements demonstrate that ciliary kinematics are not uniform across the *Paramecium* ciliary array but instead exhibit substantial regional variation in beat orientation, frequency, and metachronal wave propagation. Because the flow generated by a ciliary array is set by the density, orientation, and collective beating of its cilia, these regional differences are expected to give rise to different flows. We therefore next examined the flows generated by the ciliary array and the functions they serve.

## THE CILIARY ARRAY GENERATES STEREOTYPED FLOWS UNDERLYING SWIMMING AND FEEDING FUNCTIONS

Theoretical work has suggested that ciliates can optimize feeding efficiency by designating a specific portion of the cell surface as a “mouth,” distinct from the ciliated regions that generate swimming propulsion [4]. Having established that D1 and D2 differ architecturally and kinematically, we next asked whether these differences translate into differences in the flows the ciliary array generates and consequently into differences in the functions it performs, namely swimming and feeding.

To visualize the flow generated by the ciliary array, we performed particle image velocimetry (PIV) on immobilized cells, which continue to generate ciliary-driven flows without translating. This allowed us to measure the spatial structure of the flows with high resolution (see Methods § 1 g). The flow field consists of a pair of counter-rotating vortices on either side of the cell, with a region of high flow speed near the anterior end directed towards the oral apparatus (Fig. 3A). Flow speed is highest closest to the cell at an average of ∼ 650 *µ*m*/*s on the ventral side and about 250 *µ*m*/*s on the dorsal side, and decreases with distance from the cell, decaying to 100 *µ*m*/*s at 200 µm (Fig. 3B). This faster flow speed at the ventral side is consistent with D1 having a higher beat frequency and ciliary density (Figs. 1D, 2F).

**FIG. 3.**
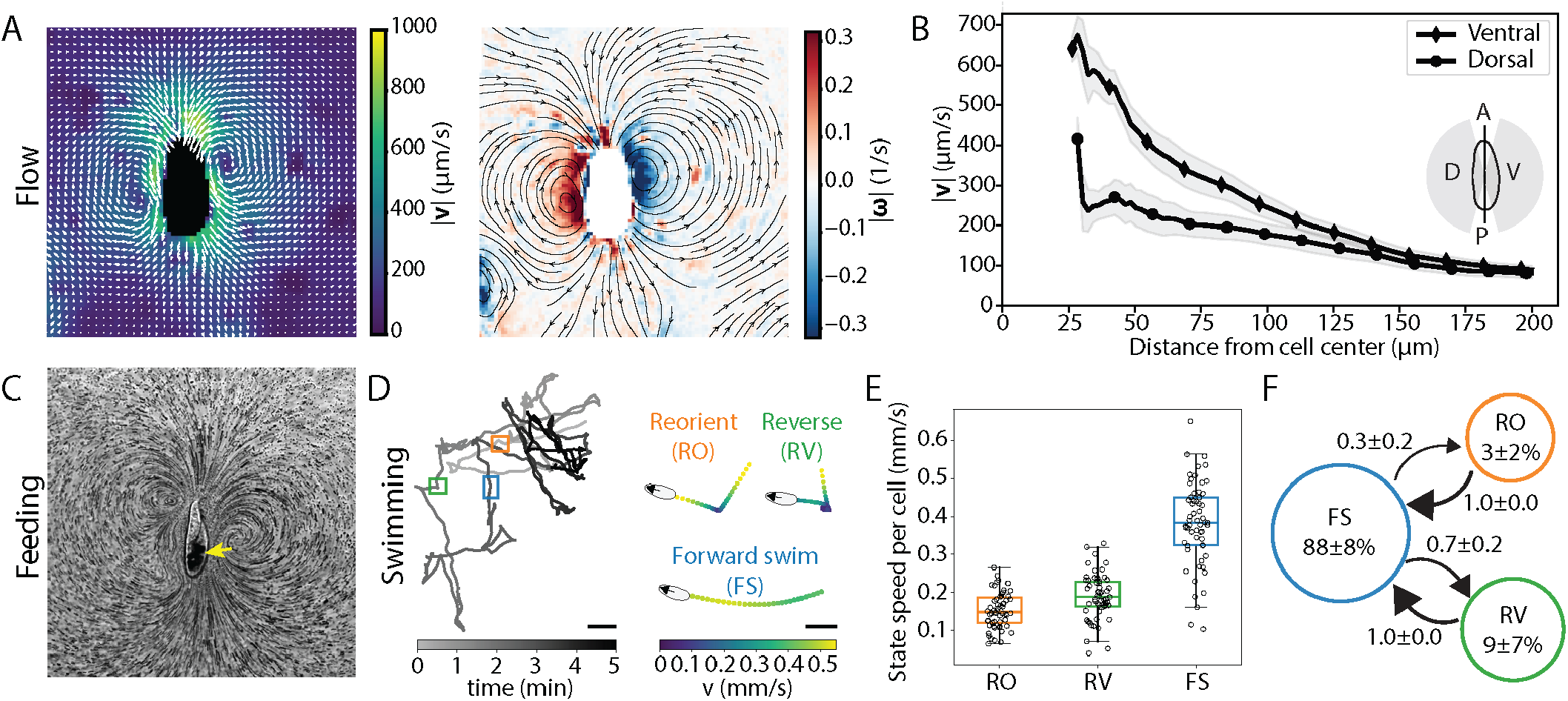
The ciliary array generates stereotyped swimming behavior and directional fluid flow. (A) Left: Representative flow field around a stationary cell immobilized with Cell-Tak, measured using fluorescent tracer beads and color-coded by flow speed. Right: corresponding vorticity field, overlaid with streamlines indicating local flow direction. Streamlines are shown only where the flow speed exceeds 5% of the maximum. (B) Mean flow velocity as a function of distance from the cell center, computed separately for the ventral and dorsal sides of stationary cells (n=4 cells). Inset: schematic defining the ventral (V) and dorsal (D) sides of the cell relative to the A–P axis, used to compute the radial profiles. Flow velocity is higher on the ventral side across the measured range. This asymmetry is consistent with the higher ciliary density and beat frequency of D1, which is located on the ventral surface (Fig. 1, 2). (C) Feeding currents around a cell in bead-containing media, visualized as a projection of a 1 s DIC movie. The yellow arrow points to food vacuoles filled with beads inside the cell body. (D) Swimming of *Paramecium* can be classified into three behavioral states: forward swimming, reorientation, and reversal. Left: Representative swimming trajectory of a single cell over 5 min, shaded by time. Colored rectangles mark example segments of each state, which are shown enlarged on the right. Right: Zoomed views of the highlighted segments, color-coded by instantaneous velocity. Duration of the segments are 1 second for reorientation and forward swimming, and 1.5 second for reversal. Scale bars: 1 mm for full trajectory and 0.1 mm for zoomed segments. (E) Swimming speed in each behavioral state. Each point represents the mean speed of one cell (n=59 cells). Mean reorientation speed: 0.15 ± 0.05 mm/s, mean reversal speed: 0.19 ± 0.06 mm/s, mean swimming speed: 0.38 ± 0.11 mm/s. (F) Probability network summarizing the observed transitions between consecutive behavioral states. The percentage of time spent in each state is shown inside each circle, and transition probabilities (mean ± SD across n=59 cells) are indicated on the connecting arrows.

Having established the structure of the flow generated by the ciliary array, we next quantitatively characterized the functional output of the array, namely feeding and swimming. To measure feeding, we exploited the well-documented tendency of *Paramecium* to ingest particles present in its environment [41, 42]. Cells were placed in a suspension of fluorescent beads (1 µm in diameter), and particle uptake was monitored by microscopy (Movie 4, see Methods § 2 e). Within seconds of exposure to the bead-containing medium, cells ingested particles, which accumulated in discrete feeding vacuoles below the oral apparatus, toward the posterior end of the cell (Fig. 3C, yellow arrow). To quantify feeding, we measured the ratio of intracellular to extracellular fluorescence intensity after 10-15 min of incubation, finding a mean ratio of 1.49 ± 0.14 (Fig. S3A), indicating substantial accumulation of ingested particles relative to the surrounding medium.

*Paramecium* swimming was first described qualitatively by Jennings from visual observations [43], who classified swimming into forward spiral motion interrupted by avoiding reactions of varying strength. Subsequent studies have measured swimming speed from microscopy videos of cells swimming in confined geometries, such as capillary tubes or microfluidic channels, over timescales of seconds[6, 44–47]. However, a complete quantitative classification of swimming into distinct gaits, and the rates of transition between them is needed to evaluate swimming function.

To address this, we imaged cells in custom-made chambers, which enable us to track freely swimming cells over minute timescales (see Methods § 1 f). We analyzed the kinematics of the cell trajectories obtained and found that each trajectory could be broken down into three gaits defined by specifying a threshold value of the deviation-angle variance (DAV, see Methods§ 2 d), a scalar parameter that captures fluctuations in instantaneous swimming direction. Our analysis identified three distinct swimming gaits, which we term forward swim (FS), reorient (RO), and reverse (RV) (Fig. 3D, Movie 3). Forward swimming is characterized by persistent, directed motion at the highest speeds. Forward swimming accounts for 88 ± 8 % of total swimming time, proceeds at 0.38 ± 0.11 mm*/*s, and occurs in bouts of highly variable duration (mean = 6.99 ± 4.43 s) (Fig. 3E,F, Fig. S3C). Reorientation is characterized by a rapid change in swimming direction with a brief drop in speed. Reorientation accounts for 3 ± 2 % of total swimming time, proceeds at 0.15 ± 0.05 mm*/*s, and occurs in brief bouts (0.50 ± 0.07 s). Reversal is characterized by a transient period of backward swimming, marked by a sign change in instantaneous velocity before a new forward direction is established. Reversal accounts for 9 ± 7 % of total swimming time, proceeds at 0.19±0.06 mm*/*s, and occurs in slightly longer bouts than reorientation (0.69 ± 0.13 s). To quantify the transitions between gaits, we calculated for each cell the probability of a transition from gait *i* to gait *j*, as done previously for the swimming states of single cells [48] (see Methods § 2 d). The resulting probability network (Fig. 3F) quantifies the first order gait transitions, while the persistence of the gaits is quantified separately by their durations (Fig. S3C). We found that turning events occur exclusively between FS bouts, so that RO and RV always return to FS. At the end of an FS bout, cells transition to RO with probability 0.3 ± 0.2 and to RV with probability 0.7 ± 0.2. *Paramecium* thus spends the vast majority of its time in persistent forward swimming, punctuated by brief RO and RV events, each identifiable by a distinct kinematic signature.

Altogether, *Paramecium* generates a structured flow field with higher velocity on the ventral surface, where cilia are denser and beat at a higher frequency (D1), and uses its ciliary array to support two distinct functional outputs. Particles are efficiently captured, reflected in the accumulation of ingested beads, and cells sustain persistent forward swimming, punctuated by brief, stochastic reorientation and reversal events. These baseline measurements of flow, feeding, and swimming provide a quantitative reference against which to assess the contribution of each ciliary domain, which we address next by selectively removing D1 or D2 cilia.

## DISTINCT CELLULAR REGIONS CARRY OUT SPECIFIC FLOW FUNCTIONS

Using these baseline measurements of flow, feeding, and swimming as a reference, we next selectively removed the cilia of either D1 or D2 to examine how each domain contributes to these functions. Deciliation of *Paramecium* cells can be triggered by chemicals such as dibucaine, ethanol, or sucrose combined with mechanical shear [5, 49–52]. We screened a range of chemical treatments to identify conditions that preferentially remove D1 or D2 cilia while preserving the remaining cilia. We found that brief ethanol exposure selectively removed D2 cilia while preserving both D1 and oral apparatus cilia (D1+OA, Fig. S4A). With longer incubation times, cilia were also lost from D1, yielding cells that retained only the cilia of the oral apparatus (OA, Fig. S4A). Sucrose treatment resulted in cells with patchy deciliation patterns. Notably, a subset of sucrose treated cells retained only the D2 and oral apparatus cilia (D2+OA, Fig. S4A). We therefore analyzed the swimming and feeding functions of cells with the desired partial ciliation patterns, to determine how each domain of the array contributes to these functions.

We first assessed the contribution of each domain to locomotion by tracking cell motility under each condition. Control cells, which were vortexed but not chemically treated, swam at 0.41 ± 0.04 mm s^*−*1^. A subset of treated cells lost only a few cilia, in both D1 and D2; we refer to this condition as Uniform (Fig. S4B). Their swimming speed (0.43 ± 0.25 mm s^*−*1^) did not differ significantly from that of control cells, although it varied more between cells. This indicates that the chemical treatments and the loss of some cilia uniformly across both regions do not detectably reduce swimming speed. D2+OA cells, which retain D2 and oral apparatus cilia but lack D1 cilia, maintained sustained forward motion at 0.30 ± 0.06 mm s^*−*1^. This speed did not differ significantly from that of uniformly ciliated cells, although it was moderately lower than that of control cells. In contrast, D1+OA cells, which retain D1 and oral apparatus cilia but lack D2 cilia, showed severely impaired locomotion, swimming slowly in spiral-like motions at 0.09 ± 0.07 mm s^*−*1^, significantly slower than uniformly ciliated cells. Cells retaining only oral apparatus cilia were effectively immotile (Fig. 4A,C, Movie 5). Statistical comparisons between all conditions are given in Supplementary Table S2. Beyond velocity, D2+OA cells also preserved the overall organization of swimming behavior. Cells lacking D1 spent most of their time in forward swimming (91 ± 6%, compared with 88 ± 8% in untreated cells; Fig. 3F) and switched from forward swimming to reverse and to reorient with probabilities of 0.7 ± 0.1 and 0.3 ± 0.1, comparable to the 0.7 ± 0.2 and 0.3 ± 0.2 of untreated cells (Fig. S4D). None of the time fractions or transition probabilities differed significantly between the two conditions (Supplementary Table S3). These results demonstrate that D2 is necessary for efficient forward propulsion and sustained locomotion, and that it is sufficient to preserve the transitions between swimming gaits.

**FIG. 4.**
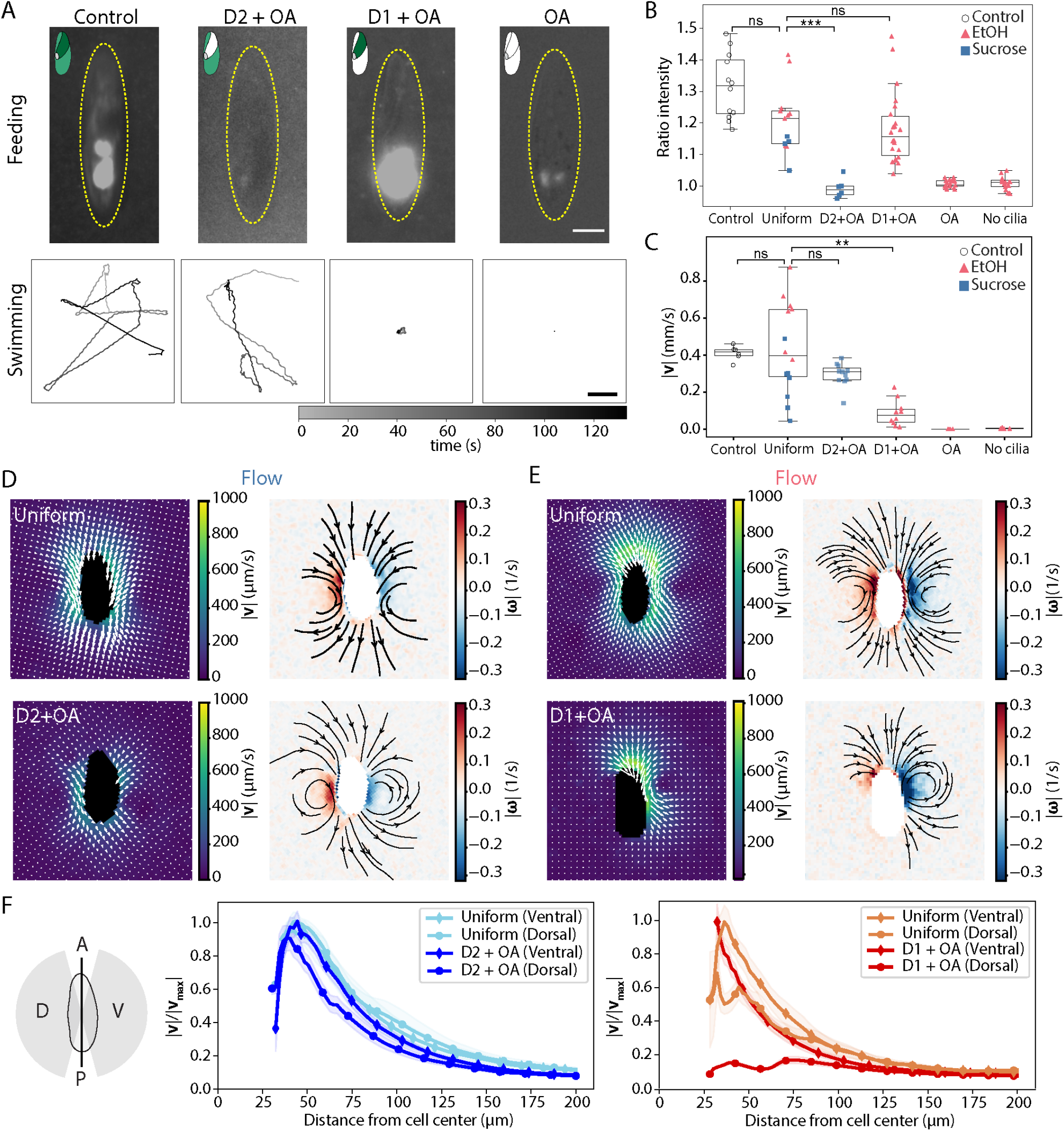
Partial deciliation reveals distinct functional roles of ciliary regions. (A) Top: Representative GFP + DIC images after feeding with fluorescent beads. The yellow dashed ellipse is the cell outline used to quantify intracellular versus extracellular fluorescence, and the schematics show in color the regions containing cilia. Scale bar: 20 µm. Bottom: Representative swimming trajectories over 130 s. Scale bar: 2 mm. (B) Feeding efficiency, quantified as the ratio of intracellular to background fluorescence intensity (n = 12, 13, 6, 22, 17, 18 cells, left to right). For comparison, feeding of cells subjected to neither vortexing nor deciliation is shown in Fig. S3A. (C) Mean swimming speed per cell (n = 6, 14, 11, 10, 5, 6 cells, left to right). In (B) and (C), circles, triangles, and squares denote control, EtOH-treated, and sucrose-treated cells, respectively. Brackets show pairwise Mann–Whitney U tests with Holm correction between control and uniformly ciliated cells, and between uniformly ciliated cells and the D2+OA and D1+OA conditions (ns, not significant; ** *p <* 0.01; *** *p <* 0.001). All comparisons are given in Supplementary Table S2. Uniform, uniformly ciliated cells; OA, cells retaining only oral apparatus cilia. (D) Flow fields around sucrose-treated cells: uniformly ciliated (top) and retaining only D2 and OA cilia (bottom). Left: flow speed. Right: vorticity with streamlines, shown where the flow speed exceeds 5% of the maximum. (E) As in (D), but for EtOH-treated cells: uniformly ciliated cell (top) and cell retaining only D1 and OA cilia (bottom). (F) Left: schematic defining the ventral (V) and dorsal (D) sides of the cell relative to the A–P axis, used to compute the radial profiles. Middle and right: Flow speed as a function of distance from the cell center, normalized to the maximum flow velocity *v*_max_, separately for the ventral and dorsal sides. Middle: sucrose-treated cells (uniformly ciliated, n=3; D2 + OA, n=5). Right: EtOH-treated cells (uniformly ciliated, n=3; D1 + OA, n=11).

In contrast to locomotion, we found that feeding is instead performed by the domain D1. Cells in the Uniform condition exhibited robust bead uptake, with a mean fluorescence ratio of 1.21 ± 0.11, not significantly different from that of control cells (1.32 ± 0.10). D1+OA cells (1.18 ± 0.11) did not differ significantly from uniformly ciliated cells, although their uptake was lower than that of control cells. In contrast, D2+OA cells showed a pronounced reduction in particle uptake, with a fluorescence ratio close to one (0.99 ± 0.03). This was significantly lower than in uniformly ciliated cells and comparable to that of fully deciliated cells or of cells retaining only oral apparatus cilia (Fig. 4A,B; Supplementary Table S2). These results demonstrate that D1 together with the oral apparatus is both necessary and sufficient for generating effective feeding flows, whereas D2 contributes minimally to particle uptake and the oral apparatus alone does not produce sufficient flow to drive particle uptake from the surroundings.

Having shown that D1 and D2 have complementary roles in swimming and feeding, we analyzed how this functional division is reflected in the contribution of each domain to the flow field of *Paramecium*. We therefore performed PIV on D2+OA cells (Fig. 4D) and D1+OA cells (Fig. 4E). In D2+OA cells, the flow vectors directed toward the mouth in the anterior region had negligible magnitudes, consistent with the impaired feeding we measured in these cells. The flow field retained a pair of counter-rotating vortices, shifted toward the posterior of the cell, and was largely symmetric between the dorsal and ventral side. Normalized radial velocity profiles confirmed this, as D2+OA cells closely matched the decay profile of uniformly ciliated cells on both the ventral and dorsal sides (Fig. 4F). In contrast, D1+OA cells showed a markedly asymmetric flow field between the dorsal and ventral sides, with large velocities directed toward the mouth near the D1 and oral apparatus regions and negligible flow on the dorsal side of the cell. Accordingly, the radial velocity profile on the ventral side decayed similarly to that of uniformly ciliated cells, whereas the dorsal velocity remained negligible at all distances from the cell (Fig. 4F).

Together, these experiments show that D1 and D2 are not only architecturally and kinematically distinct but also functionally specialized. Removing D2 severely impairs swimming while preserving feeding, and removing D1 abolishes feeding while preserving swimming. Each loss of function is accompanied by a spatially localized loss of flow in the depleted domain, indicating that D2 drives the symmetric propulsive flow required for swimming, whereas D1 generates the anterior flow directed toward the oral apparatus required for feeding. Although D1 accounts for only a small fraction of the cell surface, it is sufficient for feeding, indicating that its structural and kinematic specialization is tailored to this function. Thus, the ciliary array of *Paramecium* is not a uniform propulsive surface but a spatially partitioned system in which distinct domains independently generate the distinct flows required for swimming and feeding.

## DISCUSSION

In this work, we show that *Paramecium tetraurelia* can perform distinct flow functions with a single continuous ciliary array by partitioning its cortex into domains that differ in ciliary density, orientation, and beat kinematics. A densely ciliated, high-frequency domain anterior and to the right of the oral apparatus (D1) generates the flows required for feeding, whereas the remaining, more sparsely ciliated and slower-beating domain (D2) generates the flows required for swimming. Whereas some multicellular marine larvae can switch between feeding and swimming by modulating ciliary kinematics under neural control [8, 9], this spatial division of labor allows *Paramecium* to feed and swim simultaneously. Beyond uncovering the mechanism by which a single cell can perform multiple flow functions, our work provides, to our knowledge, the first measurements in a living cell that link ciliary architecture, together with individual and collective beat kinematics, to the resulting flows underlying distinct biological functions. These measurements provide experimental insight into how the architecture and dynamics of cilia control the functional output of a ciliary array.

Regional differences in basal body organization, biochemical modification, and beat frequency had previously been noted in *Paramecium* [14, 16–18]. However, a quantitative, global characterization of ciliary structure and kinematics that would allow local architecture to be mapped onto overall flow and function was lacking. Here, we quantify ciliary density, orientation, beat kinematics, flow, and function across scales in the same cells. By selectively removing each region, we show that the distinct regions of the ciliary array are functionally specialized. Furthermore, our results provide a quantitative description of the ciliary architecture that underlies feeding and swimming flows.

Specifically, we find that the cell is divided into two regions with distinct ciliary densities and beat frequencies. D1, located just anterior to the oral apparatus, exhibits a distinct packing geometry, higher ciliary density, and a beat frequency roughly twice that of D2, which displays lower density and slower beating. The plane of the ciliary effective stroke varies across the cell surface, following the pattern of SF orientation, which forms concentric rows around the oral apparatus anteriorly and runs parallel to the A-P axis posteriorly. This is consistent with the known role of SFs in marking cilia polarity [14, 20–24]. Despite this regional variation in beat direction, the direction of MW propagation is conserved. Consequently, the relative angle between effective stroke direction and MW propagation differs across the cell surface. We observe a dexioplectic pattern in D1, consistent with reports of dexioplectic metachrony in *P. caudatum* and *P. multimicronucleatum* [18, 37] but at odds with the purely antiplectic pattern described by earlier work specifically for this region in *P. multimicronucleatum* [36]. This discrepancy likely reflects the limitations of earlier measurements, which estimated kinematic parameters from fixed samples and relied on visual observations, although it could also be due to species-specific variation in cortical organization. In D2, we instead observe an orientation intermediate between dexioplectic and antiplectic, consistent with the dexio-antiplectic pattern described for the general body cilia in early work [36]. This partial decoupling between stroke orientation and wave direction suggests that metachronal coordination is not dictated solely by effective stroke orientation.

We find that metachronal wavelength is conserved (≈9 µm) across regions. Because cilia beat at roughly twice the frequency in D1, the wave propagates twice as fast there. These wavelengths fall within the narrow 8–30 µm range reported across diverse ciliates, including *Stentor, Tetrahymena, Spirostomum*, and *Didinium*, despite substantial variation in cell size, ciliary density, and intrinsic beat dynamics [3, 53]. This cross-species consistency, together with our observation that wavelength is conserved even between regions with different beat frequencies, supports the view that metachronal wavelength is set by physical or geometric constraints rather than tightly coupled to local beat frequency.

Understanding what sets this wavelength requires understanding how cilia coordinate into metachronal waves, which remains a central open question in *Paramecium* and in any other ciliary array. Progress has been limited by the scarcity of experimental measurements of the spatiotemporal organization of cilia that could be used to constrain theoretical models. Hydrodynamic interactions between neighboring cilia are known to play an important role [34, 54], but appear insufficient to fully explain coordination across a cell-sized array. When the anterior and posterior surfaces of a *Paramecium* cell are mechanically decoupled from fluid interactions, metachronal waves still propagate between them, pointing to an additional, cytoskeleton-mediated route of coordination through the cell’s compliant cortex [55]. This is consistent with the dense network of striated fibers and microtubule rootlets that mechanically link neighboring basal bodies [14].

Determining what sets this wavelength requires understanding how cilia coordinate into metachronal waves, which remains a central open question in *Paramecium* and in any other ciliary array. Progress has been limited by the scarcity of experimental measurements of the spatiotemporal organization of cilia that could be used to constrain theoretical models. Hydrodynamic interactions between neighboring cilia are known to play an important role [34, 54]. Therefore our measurements of cilia kinematics in immobilized cells could in principle differ from those of free swimming cells due to the boundary. Testing this would require tracking fast-swimming, rotating cells while maintaining the high spatiotemporal resolution needed to resolve the dense ciliary array, which is not currently possible. Experimental evidence suggests, however, that hydrodynamic interactions are insufficient to fully explain coordination across a cell-sized array, and that coordination relies largely on local interactions between neighboring cilia [3, 39]. In *Paramecium* when the anterior and posterior surfaces of a cell are mechanically decoupled from fluid interactions, metachronal waves still propagate between them, pointing to an additional, cytoskeleton-mediated route of coordination through the cell’s compliant cortex [55]. This is consistent with the dense network of striated fibers and microtubule rootlets that mechanically link neighboring basal bodies [14]. Therefore the ciliary beat and the metachronal waves are expected to be only weakly affected by the long-range hydrodynamic effects of the boundary, and hydrodynamics appears to play a secondary role in physiological contexts. Measuring ciliary coordination in freely swimming cells to test this directly remains an important goal for future work.

Numerical and theoretical studies have shown that metachronal waves emerge robustly from such local hydrodynamic and mechanical interactions, with dominant wavelengths set by inter-ciliary spacing and coupling strength rather than by intrinsic frequency [34, 54]. In these frameworks, changes in activity modulate beat frequency, and thus wave speed, while the spatial coordination scale remains largely fixed. Consistent with this prediction, changes in ciliary activity or fluid viscosity alter propagation velocity without substantially shifting wavelength [56]. Our results provide experimental support for this view, suggesting that regional differences in beat frequency tune wave speed to meet functional demands, while a conserved wavelength reflects the geometric and hydrodynamic constraints that set the coordination length scale of the ciliary array. Although direct theoretical work explicitly coupling metachronal wave speed to bulk flow is limited, increased beat frequency is predicted to scale with flow rate [57]. Because wave speed equals beat frequency times wavelength, and wavelength is conserved in our system, the faster waves observed in D1 are expected to generate higher local flow rates, consistent with the stronger flows we measure on the ventral side.

The origin of the twofold frequency difference between regions remains speculative. It may partly reflect regional differences in cilia density and organization, which could alter the strength of hydrodynamic and/or mechanical coupling between neighboring cilia [14], or region-specific biochemical differences in the ciliary axoneme reported elsewhere in *Paramecium* [16]. Calcium signaling, which controls the direction of ciliary beating during the avoiding reaction (see below), is a further layer of ciliary control, but whether it also shapes the coordination of cilia into metachronal waves or contributes to the differences between D1 and D2 remains to be tested.

To assess the functional relevance of the spatiotemporal differences in ciliary patterning across the cell, we combined quantitative analysis of swimming, feeding, and flow with selective regional deciliation. Swimming in *Paramecium* was first described by Jennings, who classified it based on visual observations into forward spiral motion interrupted by avoiding reactions of varying strength [43]. Our analysis recovers the categories described by Jennings as three kinematically defined gaits, forward swim, reorient, and reverse, with measurable durations and transition probabilities. Reorient and reverse events correspond to the weak and strong avoiding reactions. The avoiding reaction is triggered by a calcium-based action potential and has been modeled by coupling calcium concentration to swimming kinematics [6]. Our measurements provide a quantitative reference for such models, as well as a baseline against which to compare cells lacking either domain. Cells lacking D1 swam forward at a speed that did not differ significantly from that of uniformly ciliated cells, whereas cells lacking D2 could not sustain forward swimming and lost the flow on the dorsal side of the cell. D2 is therefore required for propulsion. Feeding showed the opposite dependence. Cells lacking D1 did not feed and lost the flow directed toward the mouth, whereas cells retaining only D1 and the oral apparatus fed at levels comparable to cells with uniform partial ciliation, so D1 is necessary for feeding and, together with the oral apparatus, sufficient for it, even though it covers only a small fraction of the cell surface. This is consistent with the earlier observation that anterior deciliation leads to a loss of feeding in *Paramecium* [50], and our results show that the anterior cilia responsible correspond to D1. Recent theoretical work predicts trade-offs between swimming efficiency and feeding performance depending on how ciliary activity is spatially distributed in unicellular organisms [4, 58]. Our findings provide experimental support for this principle, in that functional diversification requires deploying distinct ciliary spatiotemporal patterns to specific regions of the cell.

Although we have clarified the functional consequences of regional specialization, the mechanisms that generate and maintain these distinct ciliary architectures remain unknown. How D1 and D2 are patterned within a single continuous cortical array is a longstanding question. Classical work on cortical patterning in *Paramecium* showed that pre-existing cortical patterns act as templates that locally guide basal body positioning. This was demonstrated by experimental inversion of cortical domains, in which cilia with reversed beat polarity were stably inherited across divisions, indicating that patterning arises from local interactions rather than global cues [25–27]. These findings suggest that the two domains emerge through spatially restricted modulation of a self-propagating basal body pattern during cell division. Future studies targeting cortical scaffolds or basalbody-associated cytoskeletal components, for example via RNAi, could determine whether regional identity is genetically specified, mechanically stabilized, or emerges from feedback between cortical geometry and ciliary dynamics.

Perturbing this patterning genetically would also make it possible to ask how ciliary architecture relates to function. Mutants with ciliary patterning defects exist, such as *crochu*, which has an abnormal cell shape and cortical pattern [59]. How these patterning defects relate to swimming and feeding is a subject for future research.

Equally intriguing is the possibility that *Paramecium* possesses behavioral switches capable of temporally regulating each ciliary domain. If the cell can modulate the activity of D1 and D2 on demand, it would gain an additional layer of behavioral flexibility, allowing rapid adaptation to environmental changes such as threats or fluctuations in nutrient availability. Systematic investigation of swimming and feeding performance under defined environmental stimuli will be needed to test whether such temporal control exists. Demonstrating temporally regulated domain specialization would establish that functional partitioning in unicellular organisms is not purely structural but actively tunable, allowing a single cell to reconfigure its hydrodynamic output in real time.

More broadly, our work reveals general design principles for active biological surfaces that must multiplex several functions within a single continuous active surface. *Paramecium* achieves functional versatility by locally specifying distinct spatial and kinematic cilia patterns, allowing propulsion and particle uptake to occur simultaneously. These findings show how decoupling tasks through modular surface specialization can yield greater efficiency and adaptability than uniformly actuated designs, offering a blueprint for flow optimization in engineered microswimmers and biomimetic systems [60].

## METHODS

### 1. Cell preparation and imaging

#### a. Cell culture

*Paramecium tetraurelia* wild-type strain d4-2 (a gift of AnneMarie Tassin) was cultured following standard protocols [61, 62]. To obtain a homogeneous population of cells of the same age and nutritional state, cultures were synchronized through autogamy, a self-fertilization process involving nuclear reorganization that is triggered by starvation [61]. Briefly, a single cell was transferred daily into fresh medium for five days, allowing it to divide between transfers, and its progeny was then left to starve for three days. Autogamy was confirmed by Hoechst staining of the nuclei under a fluorescence microscope. The autogamous cells were washed in Dryl’s buffer (2 mM sodium citrate, 1 mM Na_2_HPO_4_, 1 mM NaH_2_PO_4_, 1.5 mM CaCl_2_) and transferred for 48 h at 27°C to standard *Paramecium* medium (wheat grass powder extract buffered with sodium phosphate and supplemented with 0.8 *µ*g/mL *β*-sitosterol), bacterized the day before with *Klebsiella pneumoniae* strain KpGe [62]. All experiments were performed on cells in the logarithmic growth phase following completion of autogamy.

#### b. Immunofluorescence staining

Cells were fixed and permeabilized using 4% paraformaldehyde (PFA) and 0.5% Triton X-100 in PHEM buffer (60mM PIPES, 25mM HEPES, 10mM EGTA, 2mM MgCl2, pH 7) for 20 minutes. After fixation, samples were washed in 0.05M glycine in PBS for 15 minutes and blocked in 0.2% bovine serum albumin (BSA) in PBS for 20 minutes. Cells were incubated with primary antibody Polyglutamate chain pAb anti-rabbit 1:2000 (Poly-E, AdipoGen AG-25B-0030-C050) and centrin anti-mouse 1:400 (Millipore ZMS1054) in antibody solution for 1 hour at room temperature. The antibody solution consists of 0.1% BSA and 0.2% Triton X-100 in PBS. After incubation with the primary antibody, the cells were washed three times (10 minutes each) in PBS with 0.2% Triton X-100. The secondary antibodies used were Alexa Fluor 568 goat anti-rabbit (Invitrogen A11004), Alexa Fluor 488-goat anti-mouse (Invitrogen A11001), and/or CenSpark to stain BBs and cilia [63], at 1:1000 dilution. Cells were incubated with the secondary antibody in antibody solution for 30-60 minutes at room temperature. Following secondary antibody staining, cells were washed twice for 10 minutes in PBS with 0.2% Triton X-100. Samples were mounted with Vectorlabs VectaShield PLUS Antifade Mounting medium (Vector Laboratories H-2000) which has 405 DAPI nuclear staining. We note that in fixed cells, a small protrusion is sometimes visible next to the oral apparatus, which is not observed in live cells and most likely results from fixation.

#### c. Ultrastructure Expansion Microscopy (U-ExM)

Expansion Microscopy was performed as in [64], with slight modifications. Cells were fixed and permeabilized in 4% PFA and 0.5% Triton X-100 in PHEM buffer for 20 minutes at room temperature. Samples were washed once with 0.05M glycine in PBS, followed by two washes with 100 mM sodium bicarbonate (pH=8.5) for 15 minutes each. Anchoring was performed by incubating cells in 0.04% Glycidyl methacrylate (GMA, SigmaAldrich 779342) in 100 mM sodium bicarbonate for 3 hours at 37°C, as previously described [65]. Prior to gelation, the anchoring solution was removed and cells were briefly incubated in monomer solution. A small drop of dense cells was allowed to sediment for approximately 10 minutes, after which excess media was removed. Gelation was carried out using a monomer solution containing 19% (wt/wt) sodium acrylate (Sigma-Aldrich 408220), 10% (wt/wt) acrylamide (Sigma-Aldrich A4058) and 0.1% (wt/wt) N,N’-methylenbisacrylamide (Sigma-Aldrich M1533) in PBS. Polymerization was initiated by adding 0.5% (wt/wt) APS (ThermoFisher 17874) and 0.5% (wt/wt) TEMED (ThermoFisher 17919) immediately before use. The gelation solution was applied to the cells and covered with a coverslip. All (pre-)gelation steps were performed at 4°C to delay polymerization and allow complete infiltration. Samples were incubated for 1 hour at 37°C in a humid chamber. Gels were denatured for 2 hours at 95°C in a preheated denaturation buffer (50 mM Tris, 200 mM NaCl, 200 mM SDS in MilliQ water, adjusted to pH 9.0 using HCl). Full expansion was achieved by incubating gels in Milli-Q water for 1 hour with gentle shaking, replacing the water every minutes. Alternatively, gels were expanded overnight at 4°C. Regions with high cell density were identified by phase-contrast microscopy. Prior to antibody staining, gels were shrunk by two 15-minute incubations in PBS. Primary antibodies were applied in antibody solution (3% BSA, 0.1% Triton X-100 in PBS) and incubated overnight (or for at least 6 hours) at 37°C with shaking. Depending on the experiment, primary antibodies included polyglutamylation (Poly-E, rabbit polyclonal, 1:400, AdipoGen AG-25B-0030-C050), anti-*α*-tubulin (ATTO 488 monoclonal, 1:400, Adipogen AG-27B-0005TD-C100), striated rootlets (rabbit polyclonal, 1:400; gifted by Anne-Marie Tassin), and/or centrin (mouse monoclonal, 1:400, Millipore, ZMS1054). Gels were washed three times for 10 minutes in PBS containing 0.1% Triton X-100 at room temperature. Secondary antibodies were incubated for at least 5 hours at 37°C with shaking. Alexa Fluor 488-conjugated goat anti-mouse IgG (Invitrogen A11001), Alexa Fluor 568-conjugated donkey anti-rabbit IgG (Invitrogen A11004), and/or Alexa Fluor 647-conjugated donkey anti-mouse IgG (Invitrogen A32787TR) were used at 1:1000 dilution in antibody solution. After secondary staining, gels were washed once in PBS for 5 minutes and incubated for 10 minutes in nuclear staining solution (NIR694 nuclear dye, 1:700, BioTracker SCT118). Gels were washed once more in PBS for 5 minutes and fully re-expanded by three successive 15-minute incubations in Milli-Q water with shaking.

For imaging, gels were attached to aminosilanized coverslips. Coverslips were sonicated sequentially in 1 M KOH (15 min), Milli-Q water (5 min) and 100% ethanol (5 min), then silanized in 2% (3-aminopropyl)triethoxysilane (APTES) in methanol for 4 min. They were rinsed in Milli-Q water and air-dried before the gels were attached.

#### d. Imaging of fixed cells

For immunofluorescence imaging used to quantify cell shape, *Paramecium* cells were imaged on an inverted microscope (Ti2, Nikon) equipped with an AX point-scanning confocal. Images were acquired using a 20x Apo LWD NA 0.95 water-immersion objective and a galvano scanner. Expanded samples were mounted on ATPES-coated coverslips (3-Aminopropyltriethoxysilane, Thermo Fisher Scientific 430941000) and enclosed within a 3mm-high acrylic chamber to maintain full immersion in water. Expanded cells were imaged on the same inverted confocal microcope using a 20x Apo LWD NA=0.95 water immersion objective and a NSPARC detector operated in super-resolution mode. Higher-resolution images of selected regions were acquired using a 40x Apo LWD NA=1.15 water-immersion objective with the NSPARC detector in super-resolution mode. These images were used to calculate the gel expansion factor.

#### e. High-speed live imaging

For live imaging of ciliary beating and metachronal waves, *Paramecium* cells were immobilized on Cell-Tak coated coverslips and glass slides using the protocol described in [35]. Imaging chambers with a height of about 180*µ*m were assembled using double-sided tape as spacers between a coated glass slide and a coated coverslip. Given a cell body thickness of approximately 40*µ*m and a ciliary length of approximately 10*µ*m, this leaves ∼130*µ*m of free fluid above the imaged cilia, about an order of magnitude larger than the ciliary length. Approximately 15 µl of standard culture media containing a high density of cells was added to each chamber. Imaging started 10 minutes after adding the cells, once cells had adhered to the coverslip. High-speed differential interference contrast (DIC) imaging was performed on an inverted Nikon Ti2 microscope equipped with a Phantom T2540 high-speed camera and a 40× Apo LWD NA 1.15 water-immersion objective. Cilia beating and metachronal waves were recorded at 1000 fps for 1-3 seconds. To visualize the direction of the effective stroke, the imaging plane was positioned approximately 10µm above the cell surface, such that ciliary tips were in focus during the effective stroke while the recovery stroke remained out of focus.

#### f. Live imaging of swimming cells

Free-swimming *Paramecium* cells were imaged in custom-built acrylic circular chambers (1 cm diameter, 1.5 mm height) mounted on a glass slide. Cells were loaded into a chamber with a syringe through an inlet. 1-3 Cells per chamber were imaged under a dissection microscope in 2D for 5 minutes at 30 fps using a Raspberry Pi HQ camera. For higher-resolution recordings of swimming behavior, DIC imaging was performed on an inverted microscope (Ti2, Nikon) with a 10x (Plan Fluor NA 1.3) or 20x (Plan Apo NA 0.8) air objective.

#### g. Flow experiments

Imaging chambers with a height of approximately 180 µm were assembled using double-sided tape as spacers between a glass slide and coverslip. Coverslips were coated with Cell-Tak to immobilize cells or left uncoated, depending on the experiment. The sides of the chamber were sealed with VALAP to prevent evaporation. To visualize flows generated by ciliary beating, 0.5 µm fluorescent beads (Polysciences 24054, diluted 1:500 in standard *Paramecium* medium) were added to the chamber as flow tracers, and DIC imaging was performed immediately thereafter on an inverted Nikon Ti2 microscope using a 20x objective, at a frame rate of 30 fps while the cells remained in frame (few seconds to 30 s).

#### h. Deciliation experiments

Partial deciliation of *Paramecium* cells was achieved using two different protocols to selectively remove cilia from specific regions. Ethanol deciliation was used to generate fully deciliated cells, cells retaining only OA cilia, or cells retaining only D1 + OA cilia [49, 50]. Cells were vortexed for 20-40 seconds in 5% ethanol in Dryl’s buffer (2 mM sodium citrate, 1 mM disodium phosphate, 1 mM monosodium phosphate, 1.5 mM calcium chloride in MilliQ water), followed by two washes in Dryl’s buffer. Sucrose deciliation was used to generate cells retaining only D2 + OA cilia [5]. Cells were vortexed for 15 s in 20% sucrose in 1mM Tris. The solution was gradually diluted with Dryl’s buffer over 5 min, followed by two washes in Dryl’s buffer. Because both treatments yield a mixture of deciliation patterns, individual cells were screened one by one by DIC microscopy prior to each experiment, using a 20x Plan Apo NA 0.8 air objective on an inverted Nikon Ti2 microscope, and only cells with the desired pattern were used. This yielded four conditions for further analysis: cells retaining only D2 and OA cilia (D2 + OA), cells retaining only D1 and OA cilia (D1 + OA), cells retaining only OA cilia, and fully deciliated cells (Fig. S4A). Cells with other patterns, such as partial loss of cilia within a domain, were excluded (Fig. S4C). Swimming behavior was imaged as described above. To verify the deciliation patterns, immunofluorescence staining and imaging was performed. In immunofluorescence images, cells lacking all cilia cannot be reliably distinguished from cells retaining cilia only in the OA. For the deciliated cells used in feeding and swimming experiments, this distinction was made by live imaging of ciliary beating in the OA.

#### i. Feeding experiments

Feeding efficiency was assessed by incubating cells with 1.0 µm fluorescent polystyrene beads (1:200 dilution, Invitrogen 2291490). Imaging was performed 10-15 minutes after bead addition using a 20x objective, acquiring GFP fluorescence with simultaneous DIC illumination to visualize both bead uptake and cell outlines.

### 2. Image analysis

#### a. Measurement of cell shape and cilia array geometry from fixed samples

Cell shape analysis. Cell shape was quantified using a custom Python script from confocal immunofluorescence images of cells stained for centrin. Centrin labels the cortical mesh that covers the whole cell surface and can therefore be used to define the cell outline. Maximum intensity projections of the centrin channel were generated and smoothed using a Gaussian filter. Cell outlines were segmented by Otsu thresholding, and properties including cell length, width, and aspect ratio were extracted using the regionprops function of the scikit-image package.

Identification of ciliary regions. Analysis of ciliary domains was performed on images of expanded *Paramecium* cells. To enable consistent measurements across cells, each cell was computationally aligned to a standardized coordinate frame. An ellipse was fitted to the cell outline in maximum-intensity projections along both the XY and YZ planes, and the cell was rotated in each plane so that the ellipse’s major axis aligned with the Y-axis. This alignment is used to standardize cell orientation for further analyses. Both rotations were applied to the full Z-stacks of the centrin channel and of the Poly-E channel, which labels polyglutamylated tubulin in basal bodies and cilia. Separate maximum projections of the two cell halves were generated. Cells were included only for further analysis when the two halves could be clearly assigned to the ventral and dorsal sides of the cell. Centrin images were used to define the total cell area, while Poly-E images were used to segment ciliary regions based on fluorescent intensity, reflecting basal body and cilia density. Fluorescence intensity is highest in the OA region, intermediate in D1, and lowest in D2. After Gaussian smoothing, the OA and D1 regions were identified on the ventral surface by intensity-based thresholding, followed by iterative erosion and dilation to remove small objects. Binary masks were smoothed and converted into polygonal regions. Regional areas were quantified using the centrin-based cell mask.

Striated fiber orientation. Striated fiber orientation was quantified from images of expanded *Paramecium* cells using an adapted Alignment by Fourier Transform (AFT) Python workflow [28]. Skeletonized maximum-intensity projections of the SF channel of either the ventral or dorsal cell surface, were generated in ImageJ and used as input for the AFT analysis. The following parameters were used: window size = 4, window overlap = 0.3, intensity threshold = 0.9, and eccentricity threshold = 0.3. Cilia density. Ciliary density was calculated from basal body spacing within and between rows. Distances were measured for both cell regions on expanded *Paramecium* cells. Intra-row spacing was measured from maximum-intensity projection Poly-E images of either the ventral or dorsal cell surface. For the D2 region, the A-P axis on the dorsal side was divided into three equal segments, and only the central third was analyzed. This restriction minimized geometric distortion arising from maximum-intensity projections of the curved cell surface, which introduce perspective-dependent spacing errors toward the edges and the anterior and posterior poles. Within this central third, the five basal body rows closest to the center of the dorsal surface were selected for each cell. For the D1 region, which was measured on the ventral surface, the five central rows of the region were selected. Intensity profiles were extracted along each row in ImageJ, smoothed, and basal body peaks were identified using the SciPy find peaks function in Python. Mean intra-row spacing was computed per row and averaged across rows for each cell. Inter-row spacing was determined from images of SFs, fibers connecting the basal bodies into rows, using a two-dimensional Fast Fourier Transform (FFT) in Python. For each region, a central area away from the cell edge was selected. The dominant spatial frequency perpendicular to the row direction was identified from the power spectrum, and inter-row spacing was calculated as the inverse of this frequency. Uncertainty was estimated from the full width at half maximum of the spectral peak. Ciliary density was calculated as the number of cilia per unit area (µm^2^) based on the mean basal body spacing withing rows and between rows:

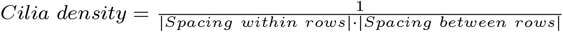

with error propagation based on the standard errors of both spacing measurements.

Expansion factor. The expansion factor of cells expanded using the expansion microscopy protocol was calculated from basal bodies diameter measurements in 40X confocal images using a custom Python script. The basal bodies (top view) were detected using Laplacian-of-Gaussian blob detection (scikit-image). Radial intensity profiles were computed from the center of each basal body, and the peak-to-peak diameter was extracted. The expansion factor was calculated by dividing the mean expanded diameter by the known basal body diameter of 200 nm [66]. Measurements were validated by comparing basal body spacing and cell lengths between expanded and non-expanded samples.

#### b. Analysis of ciliary beat direction

Ciliary beat direction was quantified from high-speed DIC microscopy recordings of immobilized cells. The focal plane was positioned approximately 10 µm above the cell surface such that only ciliary tips were visible during the effective stroke. Ciliary tip trajectories were manually traced in ImageJ over a single beat cycle using the Multi-point tool. Trajectory orientation was determined by least-squares linear regression in Python, and the resulting angle was calculated relative to the A–P axis.

#### c. Quantification of metachronal wave properties

Metachronal wave dynamics were quantified using a Python-based image analysis pipeline adapted from [3]. Image stacks were corrected for global motion using rigid-body registration of the StackReg plugin in ImageJ [67] followed by a normalization and background subtraction. First, the direction of wave propagation is determined, by placing a line of interest along the direction of propagation observed in the recording and rotating the line in 5° increments in both directions. From each line, we extract a kymograph and compute its two-dimensional autocorrelation. We then obtain the spatial wavelength from the dominant spatial peak of the autocorrelation. The wavelength measured along a line is shortest when the line is parallel to the direction of wave propagation (perpendicular to the wave fronts) since along any other orientation the line crosses neighboring wave fronts at a larger spacing. We therefore take the orientation that minimizes the wavelength as the correct direction of wave propagation (see schematic in Fig. 2C). For this identified direction, the two-dimensional autocorrelation analysis was applied. Beat frequency and wavelength were obtained from the dominant temporal and spatial peaks, and wave velocity was calculated as their product. For each cell, measurements were averaged across five parallel lines within the same region.

#### d. Analysis of cell swimming

A custom-build Python pipeline was used to detect and analyze cell swimming. Swimming videos were converted to grayscale and background subtraction was performed by computing a local median projection of 500 frames to remove static objects. Images were denoised and segmented using median filtering, thresholding, and morphological operations. Cell trajectories were extracted using the TrackPy Python package and smoothed using a Savitzky–Golay filter (window = 0.33 s, polynomial order = 2) to remove noise but preserve features like peaks or rapid changes in the trajectory. Instantaneous speed was calculated from frame-to-frame displacements. Behavioral states were classified based on directional angular variability (DAV), computed within sliding 0.5 s windows along each trajectory. DAV was quantified as the circular variance of swimming direction using the circvar function from the SciPy library. Behavioral states were assigned based on DAV thresholds: values below 0.01 were classified as swimming, values between 0.01 and 0.04 as reorientation, and values above 0.04 as reversal. Transitions were restricted such that cells could switch only between the swimming state and a turning state, but not directly between reorientation and reversal. Reorientation events immediately adjacent to reversal events were merged and classified as a single reversal. For each cell, the fraction of time spent in each behavioral state was computed, along with the distributions of state durations and swimming speeds. After classifying trajectory segments into forward swim (FS), reorient (RO) and reverse (RV) as described above, we calculated for each cell the probability of a transition from state *i* to state *j* by counting,

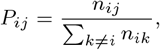

where *n*_*ij*_ is the number of transitions from state *i* to state *j* along the trajectory, excluding self-transitions. Because cells switch only between FS and a turning state, RO and RV always return to FS, and the only free probability is *P*_FS,RV_ = 1 − *P*_FS,RO_. The probability network shows the mean ± SD of *P*_*ij*_ across cells.

#### e. Feeding analysis

Feeding was quantified from images combining GFP fluorescence and DIC illumination. For each cell, an elliptical region of interest was manually drawn around the cell in ImageJ. An identical ellipse placed adjacent to the cell was used to sample background fluorescence. Mean fluorescence intensities were measured for both regions, and feeding efficiency was quantified as the ratio of intracellular to background intensity.

#### f. Statistical analysis

Statistical analysis. Differences in feeding efficiency and mean swimming speed between deciliation conditions were assessed with two-sided Mann-Whitney U tests for all pairs of conditions, with Holm correction for multiple comparisons. The time fractions of the three behavioral states and the transition probabilities out of forward swimming were compared between untreated and D2+OA cells in the same way, with Holm correction across these five comparisons. Each cell was treated as one independent observation. The p-values for all comparisons are given in Supplementary Table S2 for the deciliation conditions and in Supplementary Table S3 for the behavioral states and transitions. The sample size of each quantitative panel and the number of cells it comes from are given in Supplementary Table S1.

#### g. Analysis of flow fields

A custom-built Python pipeline was used to compute flow velocity fields from bead-tracer image sequences. Images were bandpass-filtered using a difference-of-Gaussians filter (*σ*_1_ = 2, *σ*_2_ = 12) to suppress high-frequency noise and eliminate uneven background illumination. Cell outlines were segmented in each frame using the Segment Anything Model for microscopy (micro-sam, vit-blm model) to generate binary masks. To correct for spatial drift, image stacks were aligned by calculating rigid-body transformation matrices from a cropped region centered on the cell using the StackReg Python package (pystackreg). Particle image velocimetry (PIV) was performed on the filtered image sequences using the FluidImage Python package, with a three-pass multipass scheme (initial window size ≈ 41 µm, 50% overlap, zoom factor=2 between passes) and thin-plate spline interpolation to smooth the displacement field between passes. Spurious vectors were removed based on a minimum correlation threshold (0.2) and a maximum allowed displacement difference from neighboring vectors (≈ 2.6 µm). Velocity fields were masked to exclude vectors within the segmented cell region. To account for cell motion during the recording, the transformation matrices obtained during registration were applied to each velocity field, aligning all frames to a common reference frame. Velocity fields were then averaged using a sliding window of 15 frames (0.5 s) to reduce noise while preserving flow dynamics. Local flow rotation was quantified by calculating the out-of-plane vorticity vector field (*ω*) for each frame as the curl of the in-plane flow. To establish a cellular coordinate system, cell boundaries were extracted via contour detection and fit to a parametric ellipse to determine the cell center, axis lengths, and reference orientation angle. Radial velocity profiles were extracted from 25 to 200 µm from the cell center along 12 equidistant angular trajectories, averaged within 30.0°wide wedge sectors centered on these directional axes. The dorsal and ventral sides of the cell were visually identified, and the profiles within a 120° window centered on the cell’s minor axis on each side were grouped for each cell and condition. Mean velocity profiles and standard errors of the mean across cells were then calculated as a function of radial distance from the cell center for comparative ensemble plotting.

## ACKNOWLEDGMENTS

We thank Anne-Marie Tassin for invaluable advice on *Paramecium* experimental methods and for the gift of the cell line and SF-assembly antibody. We thank Omaya Dudin and Matthew Lycas for advice on expansion microscopy. We thank Benedetta Noferi for writing the original code to measure cell shape. We thank Louise Meier for assistance with the swimming experiments. This work was supported by funding from the Swiss National Science Foundation, Project Grant 315230 215678 to G.R.S-J

## AUTHOR CONTRIBUTIONS

G.R.R.-S.J. and D.M.L. designed the research. D.M.L. performed the experiments. D.M.L., A.M.K. and G.R.R.-S.J analyzed the data. The manuscript was written by D.M.L. and G.R.R.-S.J. All authors contributed intellectually to the project. All authors edited the manuscript.

## DATA AVAILABILITY

The data that support the plots within this paper will be publicly available upon publication of the manuscript.

## CODE AVAILABILITY

The computer codes used in this paper are available from https://github.com/livingpatterns/Paramecium_cilia_functions_paper

---

**Extended Data Figure 1.**
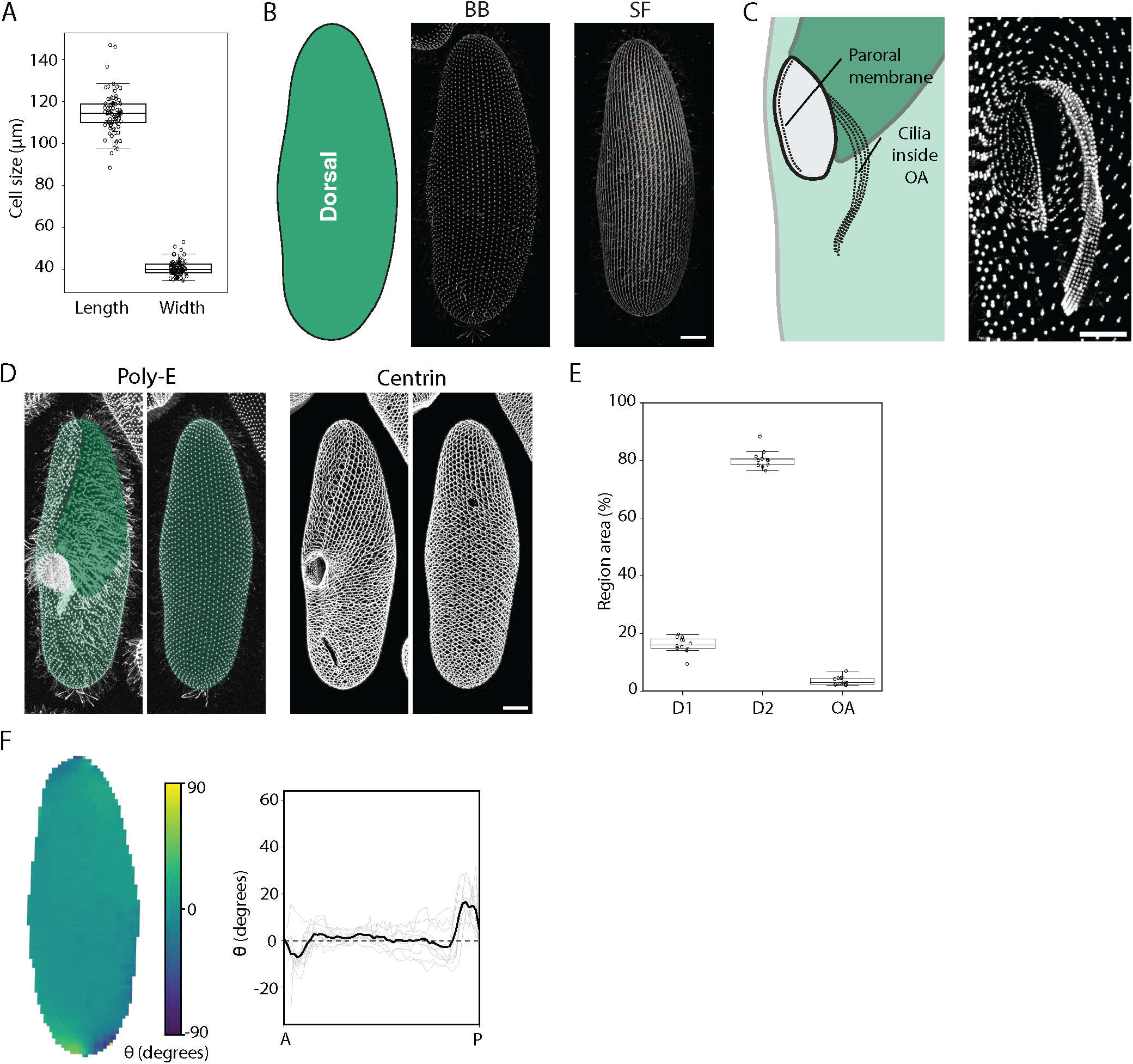
Morphological characterization of *Paramecium tetraurelia*. (A) Measurements of cell length and width. Mean cell length: 114.3 ± 9.6 µm, mean cell width: 40.3 ± 3.5 µm (n=79 cells). (B) Left: Schematic of a *Paramecium tetraurelia* showing the dorsal side of the cell (fully D2). Right: Maximum intensity projections of BBs and SF signal visualized by Expansion Microscopy (ExM). Scale bar: 10 µm. (C) Left: Schematic of the oral apparatus (OA), showing three sets of four ciliary rows inside the OA and a single row of cilia at the OA entrance forming the paroral membrane. Schematic adapted from [11]. Right: ExM maximum-intensity projection image of the ventral side of the cell showing a frontal view of the OA with basal bodies labeled by Poly-E antibody. Scale bar: 5 µm. (D) Expansion Microscopy (ExM) maximum-intensity projection images showing basal bodies and cilia stained with the Poly-E antibody (left) and the centrin mesh (right) of ventral and dorsal views of a *Paramecium tetraurelia* cell. These two channels are used to measure the region area in E. The different cell regions are marked on top of the Poly-E channel. Scale bar: 10 *µ*m. (E) Percentage of the total cell surface area occupied by each ciliary region, measured using the centrin and Poly-E channels of two-dimensional maximum-intensity projection of expanded cells. Mean percentage of area occupied by each region: D1 16.1 ± 2.8%, D2 80.4 ± 3.0%, OA 3.5 ± 1.5% (n=12 cells). (F) Left: Colormap of SF orientation angle *θ* on the dorsal cell surface, computed as in Fig. 1E, but with sectors radiating from a fixed point on the lateral edge at the cell’s A–P midpoint, rather than the oral apparatus. Right: Average SF orientation profile on the dorsal surface, computed across multiple cells as a function of angular position between the anterior and posterior ends of the cell. Gray curves represent individual cells, and the black curve indicates the population mean (n=9 cells). Dashed line marks *θ* = 0° (fibers parallel to the A-P axis).

**Extended Data Figure 2.**
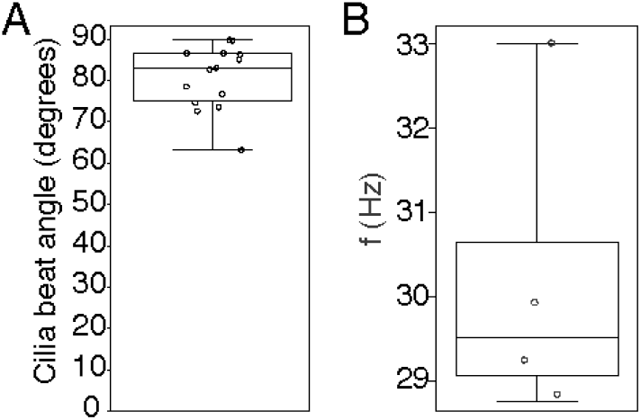
Cilia beat parameters of the paroral membrane. (A) Orientation of the paroral membrane ciliary effective stroke relative to the A-P axis. The paroral membrane is a row of cilia located at the entrance of the oral apparatus. Mean angle: 80.6 ± 7.7° (n=14 cilia trajectories from 3 cells). (B) Cilia beat frequency of the paroral membrane cilia. Mean value: 30.2 ± 2.1*Hz* (n=4 cells).

**Extended Data Figure 3.**
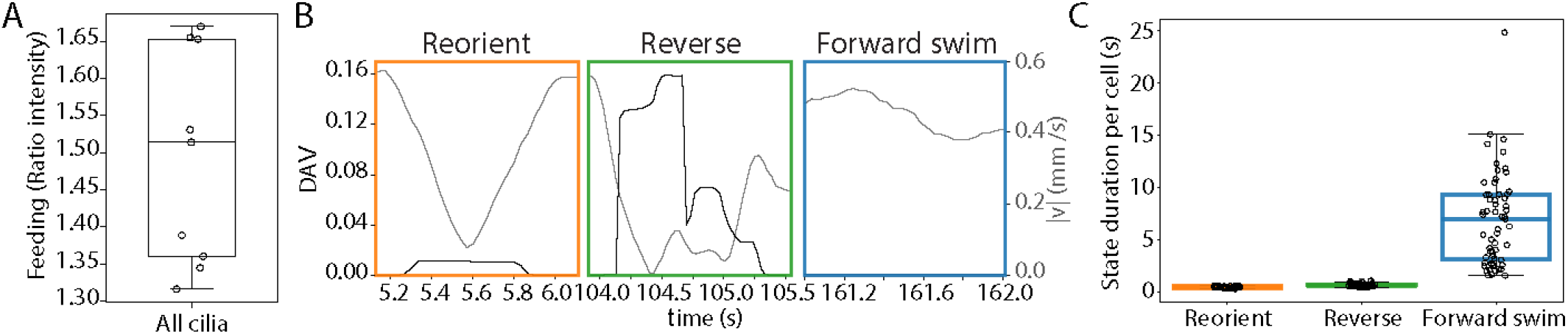
Baseline swimming and feeding characterization. (A) Feeding performance of cell not subjected to any treatment. The ratio of intracellular to extracellular fluorescence of cells not subjected to any treatment or vortexing, which therefore retain all cilia (n=9 cells). (B) Deviation angle variance (DAV, black) and instantaneous velocity (grey) for the example segments of reorientation, reversal, and forward swimming shown in Fig. 3D. Behavioral states are identified from the DAV, which measures the variability of the swimming direction along the trajectory. The DAV is near zero during forward swimming and elevated during turning events, most strongly during reversal. Velocity is not used to define the states, but drops during both reorientation and reversal. (C) Duration of each behavioral state. Each point represents the mean duration for one cell (n=59 cells). Mean ± SD: reorientation, 0.50 ± 0.07 s; reversal, 0.69 ± 0.13 s; forward swimming, 6.99 ± 4.43 s.

**Extended Data Figure 4.**
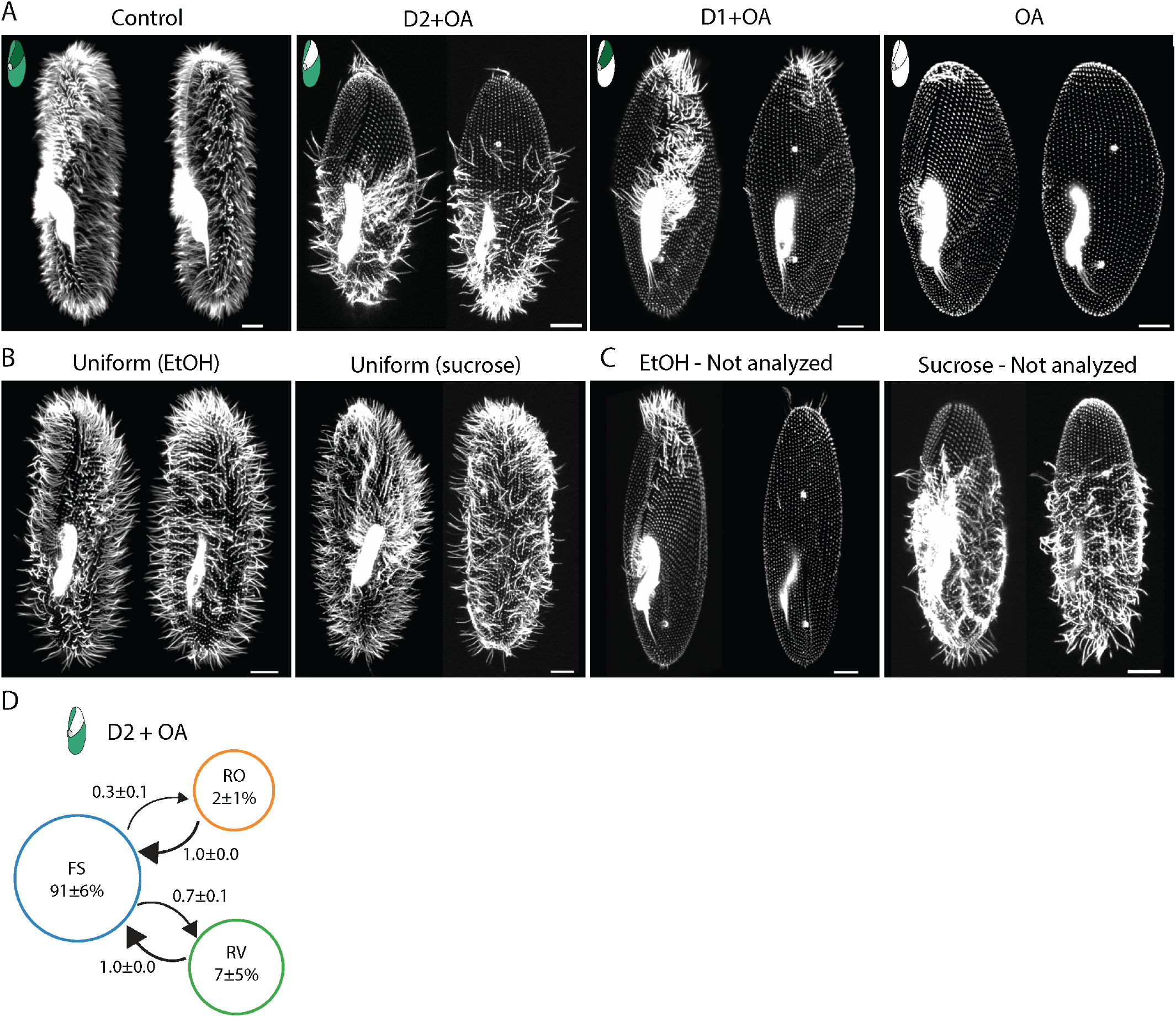
Ethanol and sucrose treatments remove cilia from distinct regions of the cell. (A) Representative immunofluorescence images of the phenotypes included in the analysis: a control cell, a cell retaining cilia only in D2 and OA, a cell retaining cilia only in D1 and OA, and a cell retaining only OA cilia. Control cells were subjected to vortexing only. Cells retaining D1 and OA cilia or only OA cilia were obtained by ethanol treatment, and cells retaining D2 and OA cilia by sucrose treatment. In ethanol-treated cells, a few cilia often remained at the cell poles. In sucrose-treated cells, part of the anterior D2 region was often also deciliated. Each cell is shown as maximum projections of its two halves (basal bodies and cilia are stained with CenSpark [63]). (B) Representative immunofluorescence images of ethanol-treated (left) and sucrose-treated (right) uniformly ciliated cells. (C) Examples of phenotypes excluded from the analysis. In the ethanol-treated cell (left), only part of the D1 region remained ciliated. Further excluded phenotypes included cells retaining D1 cilia together with part of the D2 region. In the sucrose-treated cell (right), only part of the D1 region was deciliated. Among 114 ethanol-treated cells, 9 (8%) were uniformly ciliated, 35 (31%) retained D1 and OA cilia, 3 (3%) lacked cilia or retained only OA cilia, and 67 (59%) showed other phenotypes. Among 74 sucrose-treated cells, 18 (24%) were uniformly ciliated, 7 (9%) retained only D2 and OA cilia, and 49 (66%) showed other phenotypes, including 15 cells lacking all cilia. Scale bars: 10 µm. (D) Probability network summarizing the observed transitions between consecutive behavioral states for cells retaining only D2 and OA cilia. The percentage of time spent in each state is shown inside each circle, and transition probabilities (mean ± SD across n=11 cells) are indicated on the connecting arrows.

**TABLE S1.** Sample sizes for all quantitative panels in the figures. Each entry gives the number of observations and, where these are not cells, the number of cells they come from. Unless stated otherwise, values are mean ± SD across cells, except for the stroke orientation of individual cilia (Fig. 2E and Fig. S2A), where the SD is across cilia. Groups and panels listed in one row were measured in the same cells. In Fig. 1D, D1 could not be measured in one of the cells. Representative images and trajectories are not listed. Statistical comparisons for Fig. 4B,C are given in Supplementary Table S2.

| Panel | Quantity | Group or condition | Sample size |
| --- | --- | --- | --- |
| Fig. 1D | Intra-row basal body spacing | D1 and D2 | 10 cells |
|  | Inter-row spacing | D1 and D2 | 28 cells (27 for D1) |
| Fig. 1F | SF orientation along the A-P axis | Ventral side of the cell | 10 cells |
| Fig. 2E | Orientation of the ciliary effective stroke (cells can contribute to more than one region) | Region 1 | 4 cilia in 1 cell |
|  |  | Region 2 | 17 cilia in 4 cells |
|  |  | Region 3 | 22 cilia in 5 cells |
|  |  | Region 4 | 23 cilia in 5 cells |
|  |  | Region 5 | 21 cilia in 4 cells |
|  |  | Region 6 | 18 cilia in 4 cells |
| Fig. 2F,H | Beat frequency; wave direction, wavelength, and wave velocity (all from the same cells) | D1 and D2 | 15 cells |
| Fig. 3B | Radial flow profile, stationary cells | Ventral and dorsal side of same cell | 4 cells |
| Fig. 3E | Swimming speed per behavioral state | — | 59 cells |
| Fig. 3F | Time fractions and transition probabilities | — | 59 cells |
| Fig. 4B | Feeding efficiency | Control | 12 cells |
|  |  | Uniformly ciliated | 13 cells |
|  |  | D2+OA | 6 cells |
|  |  | D1+OA | 22 cells |
|  |  | OA only | 17 cells |
|  |  | No cilia | 18 cells |
| Fig. 4C | Mean swimming speed per cell (different cells from Fig. 4B) | Control | 6 cells |
|  |  | Uniformly ciliated | 14 cells |
|  |  | D2+OA | 11 cells |
|  |  | D1+OA | 10 cells |
|  |  | OA only | 5 cells |
|  |  | No cilia | 6 cells |
| Fig. 4F | Normalized flow profile, sucrose-treated cells | Uniformly ciliated | 3 cells |
|  |  | D2+OA | 5 cells |
|  | Normalized flow profile, EtOH-treated cells | Uniformly ciliated | 3 cells |
|  |  | D1+OA | 11 cells |
| Fig. S1A | Cell length and width | — | 79 cells |
| Fig. S1E | Surface area fraction of ciliary regions | D1, D2, and OA | 12 cells |
| Fig. S1F | SF orientation along the A-P axis | Dorsal side of the cell | 9 cells |
| Fig. S2A | Stroke orientation of paroral membrane cilia | — | 14 cilia in 3 cells |
| Fig. S2B | Beat frequency of paroral membrane cilia | — | 4 cells |
| Fig. S3A | Feeding efficiency without vortexing or deciliation | — | 9 cells |
| Fig. S3C | Duration of each behavioral state | — | 59 cells |
| Fig. S4C | Deciliation phenotypes after treatment | EtOH-treated | 114 cells |
|  |  | Sucrose-treated | 74 cells |
| Fig. S4D | Time fractions and transition probabilities | D2+OA cells | 11 cells |

**TABLE S2.** Statistical comparisons between deciliation conditions (Fig. 4B,C). For feeding efficiency (A) and mean swimming speed per cell (B), descriptive statistics of each condition are followed by two-sided Mann–Whitney U tests for all 15 pairs of conditions, with Holm correction for multiple comparisons. Each cell is one observation. Sig. is based on the Holm-adjusted *p*: ns, *p* ≥ 0.05; *, *p <* 0.05; **, *p <* 0.01; ***, *p <* 0.001. *p*-values are given to four decimals, and *<*0.0001 marks values printed as 0.0000.

**A. Feeding efficiency (ratio of intracellular to background fluorescence intensity; Fig. 4B)**
| Condition | $n$ (cells) | Median | Mean | SD |
| --- | --- | --- | --- | --- |
| Control | 12 | 1.318 | 1.316 | 0.103 |
| Uniformly ciliated | 13 | 1.215 | 1.208 | 0.105 |
| D2+OA | 6 | 0.988 | 0.993 | 0.031 |
| D1+OA | 22 | 1.157 | 1.182 | 0.113 |
| OA only | 17 | 1.003 | 1.007 | 0.012 |
| No cilia | 18 | 1.012 | 1.008 | 0.020 |

| Group 1 | Group 2 | $p$ (raw) | $p$ (Holm) | Sig. |
| --- | --- | --- | --- | --- |
| Control | Uniformly ciliated | 0.0240 | 0.1199 | ns |
| Control | D2+OA | 0.0001 | 0.0008 | *** |
| Control | D1+OA | 0.0014 | 0.0086 | ** |
| Control | OA only | <0.0001 | 0.0001 | *** |
| Control | No cilia | <0.0001 | 0.0001 | *** |
| Uniformly ciliated | D2+OA | 0.0001 | 0.0006 | *** |
| Uniformly ciliated | D1+OA | 0.4029 | 0.8058 | ns |
| Uniformly ciliated | OA only | <0.0001 | <0.0001 | *** |
| Uniformly ciliated | No cilia | <0.0001 | <0.0001 | *** |
| D2+OA | D1+OA | <0.0001 | 0.0001 | *** |
| D2+OA | OA only | 0.0736 | 0.2946 | ns |
| D2+OA | No cilia | 0.1036 | 0.3107 | ns |
| D1+OA | OA only | <0.0001 | <0.0001 | *** |
| D1+OA | No cilia | <0.0001 | <0.0001 | *** |
| OA only | No cilia | 0.7043 | 0.8058 | ns |

**B. Mean swimming speed per cell (mm/s; Fig. 4C)**
| Condition | $n$ (cells) | Median | Mean | SD |
| --- | --- | --- | --- | --- |
| Control | 6 | 0.418 | 0.412 | 0.039 |
| Uniformly ciliated | 14 | 0.397 | 0.432 | 0.249 |
| D2+OA | 11 | 0.310 | 0.295 | 0.064 |
| D1+OA | 10 | 0.074 | 0.087 | 0.070 |
| OA only | 5 | 0.001 | 0.001 | 0.000 |
| No cilia | 6 | 0.003 | 0.005 | 0.003 |

| Group 1 | Group 2 | $p$ (raw) | $p$ (Holm) | Sig. |
| --- | --- | --- | --- | --- |
| Control | Uniformly ciliated | 0.9044 | 0.9044 | ns |
| Control | D2+OA | 0.0006 | 0.0049 | ** |
| Control | D1+OA | 0.0002 | 0.0027 | ** |
| Control | OA only | 0.0043 | 0.0173 | * |
| Control | No cilia | 0.0022 | 0.0108 | * |
| Uniformly ciliated | D2+OA | 0.1798 | 0.3597 | ns |
| Uniformly ciliated | D1+OA | 0.0006 | 0.0049 | ** |
| Uniformly ciliated | OA only | 0.0002 | 0.0023 | ** |
| Uniformly ciliated | No cilia | 0.0001 | 0.0008 | *** |
| D2+OA | D1+OA | 0.0002 | 0.0026 | ** |
| D2+OA | OA only | 0.0005 | 0.0041 | ** |
| D2+OA | No cilia | 0.0002 | 0.0023 | ** |
| D1+OA | OA only | 0.0007 | 0.0049 | ** |
| D1+OA | No cilia | 0.0002 | 0.0027 | ** |
| OA only | No cilia | 0.0043 | 0.0173 | * |

**TABLE S3.** Comparison of behavioral states and transitions between untreated and D2+OA cells. Values are mean ± SD across cells (*n* = 59 untreated cells, *n* = 11 D2+OA cells; Fig. 3F and Fig. S4D). Time fractions are given as the percentage of the recorded time spent in each state. Transition probabilities are the probabilities that a forward swimming bout ends in the indicated state, estimated per cell. Groups were compared with two-sided Mann–Whitney U tests and *p*-values were corrected across the five comparisons with the Holm method. The two transition probabilities out of FS sum to one in every cell and therefore yield the same *p*-value. ns, not significant.

| Quantity | Untreated | D2+OA | $p$ | $p$ (Holm) | Significance |
| --- | --- | --- | --- | --- | --- |
| Time fraction, FS (%) | $87.7 \pm 8.4$ | $90.8 \pm 5.6$ | 0.27 | 1.00 | ns |
| Time fraction, RO (%) | $2.8 \pm 2.3$ | $1.7 \pm 0.7$ | 0.34 | 1.00 | ns |
| Time fraction, RV (%) | $9.5 \pm 7.0$ | $7.5 \pm 5.3$ | 0.38 | 1.00 | ns |
| Transition probability, FS to RV | $0.70 \pm 0.17$ | $0.68 \pm 0.14$ | 0.56 | 1.00 | ns |
| Transition probability, FS to RO | $0.30 \pm 0.17$ | $0.32 \pm 0.14$ | 0.56 | 1.00 | ns |

Movie 1. **Ciliary effective stroke orientation in distinct ciliary regions**. High-speed DIC recordings of immobilized cells showing the ciliary effective stroke used to determine beat direction in different regions of *Paramecium tetraurelia*. The first segment shows the ventral cell surface with cilia tips in focus in the D1 region. The second segment shows the dorsal cell surface, capturing ciliary beating in D2. The last segment shows the beating of the paroral membrane cilia at the opening of the OA. Recordings were required using DIC microscopy with a 40x objective at 1000 fps and the video is slowed down 20 times for visualization. Scale bars: 10 µm

Movie 2. **Metachronal wave propagation on the cell surface**. High-speed DIC recording of showing metachronal wave propagation on the ventral surface of an immobilized *Paramecium tetraurelia* cell, corresponding to the cell shown in Fig. 2D. Imaging was performed using a 40x objective at 1000 fps and the video is slowed down 20 times for visualization. Scale bar: 10 µm

Movie 3. **Swimming behaviors of *Paramecium tetraurelia***. Free-swimming *Paramecium tetraurelia* cells shown in real time, followed by slowed segments illustrating transitions between forward swimming and the two different turning states, reorientation and reversals. Video was acquired using DIC microscopy with a 10x objective at approximately 45 fps.

Movie 4. **Particle uptake from surrounding media**. Immobilized cell incubated in medium containing 1 µm fluorescent beads. Forming food vacuoles inside *Paramecium* appear as dark spots inside the cell. The video was acquired using DIC microscopy with a 20x objective at approximately 30 fps. The fluorescent beads are therefore not visualized by their fluorescence in this recording.

Movie 5. **Swimming behavior following partial deciliation**. Free-swimming *Paramecium tetraurelia* cells subjected to partial deciliation. Each segment indicates the remaining ciliary regions. Cells retaining only cilia in the D1 and OA regions exhibit strongly impaired swimming. Videos were acquired using DIC microscopy with 10x or 20x objectives and are shown in real time.

Movie 6. **Recovery of swimming behavior following deciliation**. Free-swimming *Paramecium tetraurelia* cells incubated in Dryll’s buffer following ethanol-induced deciliation. The first segment shows cells 10 minutes after deciliation, when most cells retain only OA cilia or OA cilia and D1 cilia. The second segment shows the same cells 5 hours after deciliation, during partial cilia regrowth. The final segment shows the same cells after overnight (ON) recovery, when ciliation and swimming behavior are restored. Videos were acquired using a Raspberry Pi camera mounted to a dissection microscope at 30 fps.

